# An envelope stress response governs long-chain fatty acid metabolism via a small RNA to maintain redox homeostasis in *Escherichia coli*

**DOI:** 10.1101/2024.10.18.618624

**Authors:** Megha Shrivastava, Manmehar Kaur, Liz Maria Luke, Richa Ashok Kakkar, Deeptodeep Roy, Shivam Singla, Vanshika Sharma, Gaurav Sharma, Rachna Chaba

## Abstract

Long-chain fatty acids (LCFAs) are a tremendous source of energy for several bacteria but are complex to use because they induce redox stress. We previously showed that LCFA degradation impedes oxidative protein folding in the *Escherichia coli* envelope, an issue that arises due to the insufficiency of ubiquinone, a lipid-soluble electron carrier in the electron transport chain (ETC). To maintain redox homeostasis, *E. coli* activates the CpxAR two-component system; however, the nature of feedback imparted by this envelope stress response (ESR) remained unknown. Here, we show that contrary to the well-recognized remedial mode of Cpx restoring envelope integrity by upregulating protein quality control factors, in LCFA-grown cells, it uses a preventive measure to maintain homeostasis. Cpx increases ubiquinone availability for oxidative protein folding by suppressing LCFA metabolism and directly increasing ubiquinone levels. Further, rather than using its conventional mode of imparting regulation via CpxR working as a transcriptional regulator, during LCFA metabolism, Cpx mainly uses its non-coding arm to counteract envelope redox stress. The Cpx-regulated small RNA CpxQ represses *fad* genes involved in LCFA transport and β-oxidation, downregulates components of glyoxylate shunt, gluconeogenesis, and ETC, and increases ubiquinone content. Corroborating with its role in repressing LCFA metabolism and maintaining redox homeostasis, CpxQ overexpression impairs growth of *E. coli* in LCFAs and CpxQ deletion renders LCFA-grown *E. coli* hypersensitive to a thiol agent. Our foremost work studying the interconnection between LCFA metabolism, redox stress, and ESR in *E. coli* provides a rationale for investigating similar networks in other LCFA-utilizing bacteria.

**Significance Statement:** Long-chain fatty acids (LCFAs) are energy-rich nutrients for *Escherichia coli*; however, their utilization hampers disulfide bond (DSB) formation in secreted proteins, an essential process that occurs in the envelope compartment. Here, we show that an envelope stress response manipulates LCFA metabolism in *E. coli* and uses a small RNA (sRNA) to restore homeostasis. Several bacteria with a huge impact on human health use host-derived LCFAs during infection. Because many virulence factors require DSB formation, the present study offers a basis to examine whether sRNAs play a role in governing envelope redox balance during LCFA metabolism in pathogens. The sRNA-mediated control is likely an ideal strategy both for rapid response to and quick recovery from LCFA-induced stress.

## Introduction

Long-chain fatty acids (LCFAs) are carboxylic acids with a linear aliphatic chain of 12–20 carbon atoms. As energy-rich nutrients, the role of LCFAs in supporting bacterial survival and infectivity is well-exemplified by the upregulation of LCFA degradation pathway in *Mycobacterium tuberculosis*, *Pseudomonas aeruginosa*, *Salmonella* Typhimurium, and *Vibrio cholerae* during infection and the reduced virulence of their mutants defective in LCFA utilization (1–8). Most of our knowledge of the LCFA catabolic pathway is derived from studies in *E. coli*. Briefly, the Fad (fatty <u>a</u>cid <u>d</u>egradation) proteins transport and activate exogenous LCFAs to acyl_(n)_-CoA, and via β-oxidation degrade them to acetyl-CoA, which then enters the tricarboxylic acid (TCA) and glyoxylate cycles (9, 10). During aerobic metabolism, reduced cofactors produced in the β-oxidation and TCA cycle are oxidized in the electron transport chain (ETC) by respiratory dehydrogenases, and the electrons are transferred to ubiquinone, a lipid-soluble electron carrier. The reduced form of ubiquinone, ubiquinol, in turn, donates electrons to molecular oxygen via terminal oxidases. During electron transfer through ETC, proton motive force is generated, which is then used by ATP synthase to drive ATP synthesis (Fig S1) (11). Because LCFAs are non-fermentable carbon sources, TCA and glyoxylate cycles, and gluconeogenesis are critical for the generation of metabolite precursors for growth, and a functional ETC is crucial for energy generation by oxidative phosphorylation (9, 12, 13). In *E. coli*, LCFA metabolism is largely known to be regulated at a transcriptional level. The direct transcriptional regulation on aerobic *fad* genes is mainly via the three systems: (i) repression by FadR, relieved by binding of acyl_(n)_-CoA, (ii) repression by ArcBA two-component system under anaerobic conditions, and (iii) activation by the cyclic AMP-cyclic AMP receptor protein (cAMP-CRP) complex (9, 10). Besides *fad* genes, ArcBA and cAMP-CRP also directly regulate several components of TCA and glyoxylate cycles, gluconeogenesis, and ETC (12, 14–18). Contrary to the transcriptional control, relatively little is known about the post-transcriptional regulation of *fad* genes by small RNAs (sRNAs). Amongst Fad components, only FadL, the outer membrane transporter of LCFAs, and FadE, the acyl-CoA dehydrogenase of β-oxidation are known to be regulated by sRNAs. FadL is repressed by RybB, whereas FadE is activated by DsrA and ArcZ (19, 20). The sRNAs, FnrS, GcvB, GlnZ, RybB, RyhB, SgrS, and Spf regulate a few components of the TCA cycle, gluconeogenesis, and ETC (21–25).

Although LCFA metabolism has been studied in great detail in *E. coli* since the 1960s, its interconnection with other cellular processes remained largely unknown. We recently uncovered a novel link between LCFA metabolism and the redox state of the envelope, a multilayered compartment surrounding the cytoplasm, which protects the cell from environmental stresses and is the site for myriad functions critical for growth and viability. Briefly, a comparative analysis of our dataset obtained from a high-throughput genetic screen of the *E. coli* gene deletion library on oleate (C18:1 cis-9 LCFA) with similar datasets available for other non-fermentable carbon sources revealed a maximal requirement of ubiquinone for growth in LCFAs. We showed that the increased requirement of ubiquinone in LCFAs is to rapidly transfer a large number of electrons derived from its metabolism and thus decrease the formation of reactive oxygen species (13). Besides shuttling electrons derived from metabolism, ubiquinone plays a pivotal role in re-oxidizing the disulfide bond (DSB)-forming machinery (DsbA-DsbB), which forms DSBs in secreted proteins in the oxidizing environment of the periplasm (Fig S1) (26, 27). We showed that in LCFA-utilizing cells, there is a gradual increase in load on the DSB-forming machinery with increasing cell density, with a maximal load at entry into the stationary phase. Importantly, the hallmarks of inadequate DSB formation are prevented in cells exogenously provided with ubiquinone. Taken together, our work suggested that an increased electron flow in the ETC during LCFA metabolism titrates ubiquinone, limiting its availability for DSB formation, thereby compromising envelope redox homeostasis (Fig S1) (28).

In *E. coli*, ∼300 secreted proteins are predicted to have DSBs (29). Several of these involved in diverse biological processes are reported to depend on DsbA-DsbB for correct folding (27, 30, 31). Therefore, to ensure cellular integrity, bacteria must monitor the envelope redox status and elicit an appropriate response when disturbances occur. Amongst the five committed envelope stress response (ESR) systems in *E. coli*, which sense problems in the envelope and reprogram transcription to counteract stress, we identified the CpxAR two-component system as the major ESR activated by LCFAs in the stationary phase (Fig S1) (28). When cells encounter envelope stress, the inner-membrane sensor kinase CpxA, auto-phosphorylates and then phosphorylates the cytoplasmic response regulator CpxR, which, in turn, directs the transcription of its target genes involved in mitigating stress (32, 33). A well-documented mechanism by which Cpx restores envelope homeostasis is by upregulating envelope-localized protein folding and degrading factors that manage damaged proteins (32, 34). Because compromised DSB formation in LCFA-grown cells will result in the accumulation of unfolded/misfolded proteins, Cpx might restore envelope homeostasis by repairing/clearing out these proteins. However, our previous observation that ubiquinone accumulates in LCFA-utilizing cells in a CpxR-dependent manner suggested that an effective mechanism for restoring envelope redox homeostasis would be to facilitate DSB formation by increasing the oxidizing power of the ETC (28). Besides directly upregulating ubiquinone, it is plausible that Cpx also increases ubiquinone availability for DSB formation by downregulating components of the LCFA metabolic pathway that feed electrons into the ETC, an aspect that has not been examined.

Here, we investigated the global regulation mediated by Cpx during LCFA metabolism by performing RNA-Sequencing (RNA-Seq) of wild-type (WT) *E. coli* and its isogenic Δ*cpxR* strain grown in LCFAs. We found that CpxR downregulates several processes critical for the generation of reduced cofactors, metabolites, and energy during LCFA metabolism, and uses CpxQ sRNA to mediate its effect-CpxQ downregulates various *fad* genes involved in β-oxidation likely by increasing the pool of apo-FadR (not bound to acyl_(n)_-CoA), a consequence of CpxQ-mediated repression of the outer-membrane LCFA transporter, FadL, and also downregulates few components of the glyoxylate shunt, gluconeogenesis, and ETC. We further probed the regulation of ubiquinone by Cpx in LCFA-grown cells, and here again, observed that CpxR increases ubiquinone levels post-transcriptionally via sRNAs-CpxQ stabilizes OmrA in LCFA-grown cells and both sRNAs are required for ubiquinone accumulation. Corroborating with its role as a repressor of LCFA metabolism, CpxQ overexpression prevented growth of *E. coli* in LCFAs, and supporting its role in maintaining redox homeostasis, CpxQ deletion rendered LCFA-grown *E. coli* hypersensitive to a thiol agent. Collectively, our work identified CpxR as a global regulator that manages envelope redox stress in LCFA-grown cells by downregulating LCFA metabolism and increasing ubiquinone content mainly via the sRNA CpxQ.

## Results

### Transcriptomics reveals Cpx as a global regulator of LCFA metabolism

To enable a holistic view of Cpx regulation during LCFA metabolism, we performed RNA-Seq for *E. coli* BW25113 (henceforth referred to as WT) and its isogenic Δ*cpxR* strains cultured in buffered tryptone broth (TBK) supplemented with either oleate (TBK-Ole) or with detergent Brij-58 used for solubilizing oleate (TBK-Brij) in the stationary phase. Because we previously showed that the Cpx pathway is activated in LCFA-grown *E. coli* MG1655 in the stationary phase (28), here, using a chromosomal *cpxP*-*lacZ* transcriptional reporter (*cpxP* is a *bona fide* Cpx regulon member (35)), we first validated Cpx activation in the stationary phase in LCFA-grown *E. coli* BW25113 as well. Whereas the expression of *cpxP*-*lacZ* was similar in cells grown in TBK-Brij and TBK-Ole in the exponential phase (time point T1), there was a ∼4-fold increase in *cpxP* expression in TBK-Ole compared to TBK-Brij in the stationary phase (time point T2) (Figs S2A and B).

Using reads obtained from RNA-Seq, we performed differential expression analysis, followed by gene-set enrichment analysis (GSEA) to identify pathways that are enriched in TBK-Ole compared to TBK-Brij and that are regulated by CpxR (Supplementary datasets S1 and S2, Figs S3 and S4). Before probing deeper into these datasets, we validated the robustness of our RNA-Seq and our statistical approach. Notably, GSEA for WT cultured in TBK-Brij and TBK-Ole highlighted enrichment of LCFA transport and β-oxidation (FDR q value: 0%), critical for the utilization of LCFAs as an energy source (13). Additionally, TCA cycle and glyoxylate shunt (FDR q value <0.4%), gluconeogenesis (FDR q value: 7.7%) and multiple pathways for electron transfer activity (FDR q value <6%), pathways critical for the generation of reduced cofactors, metabolites, and energy during LCFA metabolism (13), were also significantly enriched. In differential expression analysis, the components of the above pathways were mainly upregulated in TBK-Ole (Figs 1A and S3, Supplementary datasets S1 and S2, and Table S1).

**Figure 1.**
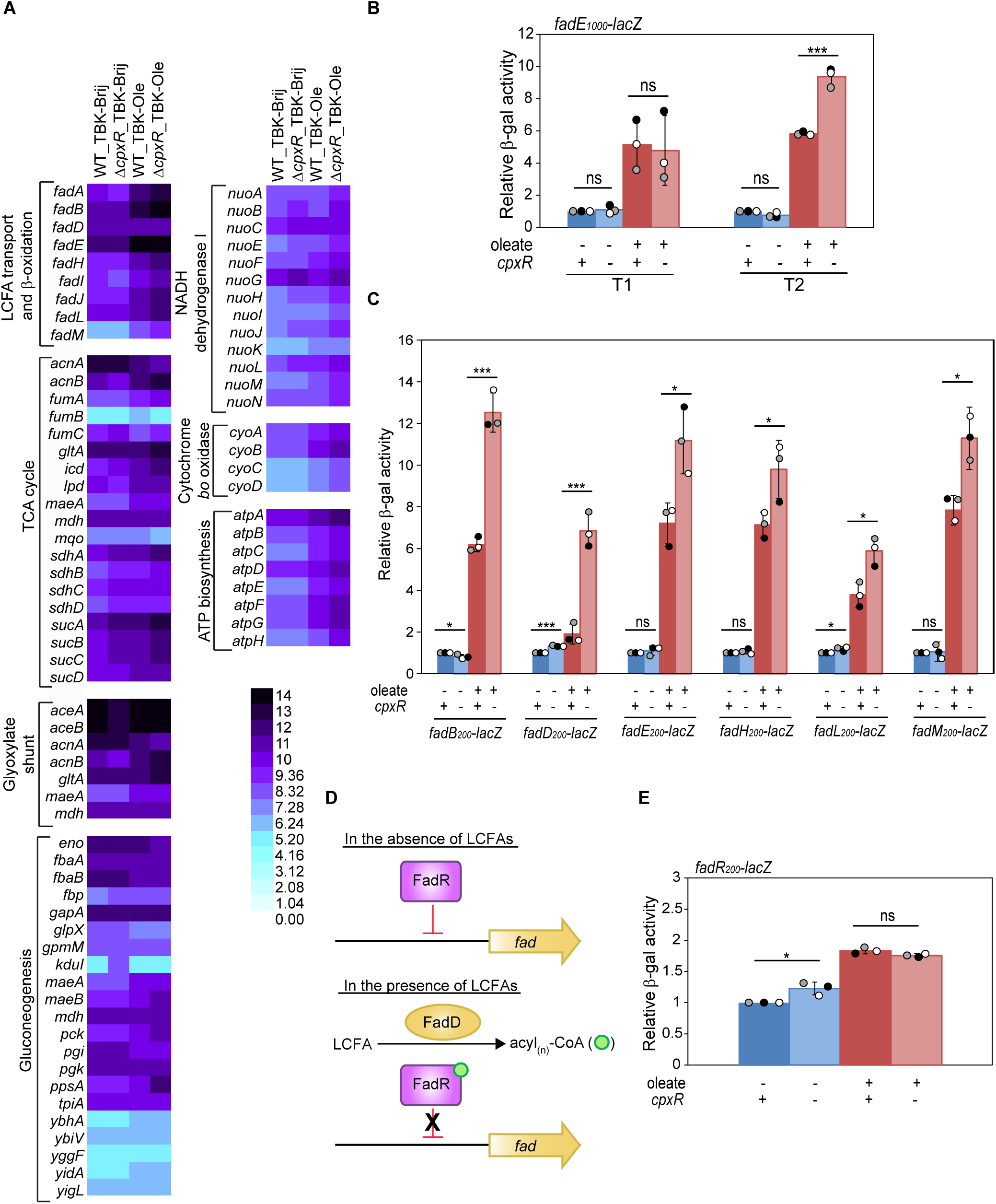
CpxR downregulates LCFA metabolism in the stationary phase. *A)* Heatmap for comparison of gene expression in four different samples WT_TBK-Brij, Δ*cpxR*_TBK-Brij, WT_TBK-Ole, and Δ*cpxR*_TBK-Ole, for various LCFA-related pathways enriched in GSEA, using normalized values provided in Table S1. The color scale represents the normalized counts. The values represented in the heatmap have been proportionally scaled down by the heatmap Illustrator for ease of representation and visualization. *B)* CpxR represses *fadE* in oleate-grown cells only in the stationary phase. Strains carrying *fadE_1000_*-*lacZ* transcriptional reporter were grown either in TBK-Brij or TBK-Ole. Cultures were harvested in the exponential and stationary phases of growth. β-gal activity was measured, and data were normalized to the β-gal activity of WT grown in TBK-Brij at respective phases. Data represent the average (± SD) of three independent experiments. The average β-gal activity (in Miller units) of WT in TBK-Brij at exponential and stationary phases was 48±21 and 92±27, respectively. *C)* Several *fad* genes are repressed by CpxR in oleate-grown cells. Strains carrying chromosomal fusion of *lacZ* with the promoter of various *fad* genes were grown either in TBK-Brij or TBK-Ole. Cultures were harvested in the stationary phase, and β-gal activity was measured. For each reporter fusion, data were normalized to the β-gal activity of WT in TBK-Brij and represent the average (± SD) of three independent experiments. The average β-gal activity (in Miller units) of various WT reporter strains in TBK-Brij was: *fadB_200_-lacZ* (62±10), *fadD_200_-lacZ* (222±1), *fadE_200_-lacZ* (85±8), *fadH_200_-lacZ* (88±16), *fadL_200_-lacZ* (76±10), and *fadM_200_-lacZ* (48±9). *D)* Regulation of *fad* genes by FadR. In the absence of LCFAs, FadR transcriptionally represses the expression of *fad* genes. In the presence of LCFAs, the esterified product of FadD, acyl_(n)_-CoA, binds FadR and releases it from the operator of *fad* genes (10). *E)* CpxR does not regulate FadR transcriptionally. Strains carrying *fadR_200_-lacZ* transcriptional reporter were grown either in TBK-Brij or TBK-Ole. Cultures were harvested in the stationary phase, and β-gal activity was measured. Data were normalized to the β-gal activity of WT in TBK-Brij. Data represent the average (± SD) of three independent experiments. The average β-gal activity (in Miller units) of WT in TBK-Brij was 34±1. For panels B, C, and E, the p-values were calculated using the unpaired two-tailed Student’s t test (***, P<0.001; **, P<0.01; *, P<0.05; ns, P>0.05).

A comparison of GSEA datasets of WT and Δ*cpxR* in TBK-Ole highlighted CpxR-dependent enrichment of almost all pathways involved in LCFA metabolism; LCFA transport and β-oxidation (FDR q value: 0.2%), TCA cycle and glyoxylate shunt (FDR q value <4%), gluconeogenesis (FDR q value: 0.2%), multiple pathways for electron transfer activity (FDR q value <3%), and ATP biosynthesis (FDR q value: 0%). In particular, in differential expression analysis, components of these pathways were mainly negatively regulated by CpxR (Figs 1A and S4B, Supplementary datasets S1 and S2, and Table S1). These data suggested CpxR as a global repressor of LCFA metabolism and encouraged us to gain further insights into its regulation.

### Cpx downregulates *fad* genes involved in LCFA transport and β-oxidation

Cpx was shown to modulate the TCA cycle and ETC in *E. coli* K-12 and enteropathogenic *E. coli* (EPEC), and to directly repress the *nuo* and *cyo* operons in EPEC (34, 36, 37); however, its effects during growth in LCFAs were unknown. We, therefore, performed an in-depth investigation of Cpx-mediated regulation of *fad* genes, specific to LCFA utilization. Using a chromosomal transcriptional reporter where *lacZ* was fused downstream of the ∼1000 bp *cis*-acting element of *fadE* (*fadE*_1000_-*lacZ*), we previously showed that FadE is induced in LCFA-grown cells (28). Here, using this reporter, we validated that *fadE* is negatively regulated by Cpx in LCFA-grown cells; whereas the expression of *fadE*_1000_-*lacZ* was similar in WT and Δ*cpxR* strains grown in TBK-Brij, in TBK-Ole there was a ∼1.6-fold increase in its expression in Δ*cpxR* compared to WT (Fig 1B, see T2), and this increase was observed only in the stationary phase where Cpx response is activated (compare Figs 1B and S2B). The truncation analysis of the *cis*-acting element of *fadE* (*fadE*_400_-*lacZ* and *fadE*_200_-*lacZ*) showed that the site required for Cpx regulation lies in the ∼200 bp region upstream of the translational start site (Fig S5A); similar to *fadE*_1000_-*lacZ*, the expression of *fadE*_200_-*lacZ* also increased ∼2-fold in LCFA-utilizing Δ*cpxR* only in the stationary phase (Figs 1B and S5B). Through transcriptional reporter assays, we confirmed that several other *fad* genes (*fadB*_200_-*lacZ*, *fadD*_200_- *lacZ*, *fadH*_200_-*lacZ*, *fadL*_200_-*lacZ*, and *fadM*_200_-*lacZ*) are also repressed by Cpx in the stationary phase (∼1.5 to 3.5-fold) (Fig 1C).

### Cpx downregulates *fad* genes post-transcriptionally via CpxQ

CpxR regulates multiple *fad* genes involved in LCFA transport and β-oxidation (Fig 1C). Except for *fadB* and *fadA*, which are part of an operon, all other *fad* genes are scattered on the chromosome (10), raising the possibility that CpxR modulates *fad* genes via a common regulator. A likely candidate is FadR, the LCFA-specific direct transcriptional repressor of *fad* genes (Fig 1D) (10); CpxR might transcriptionally induce *fadR* increasing FadR protein levels such that acyl_(n)_-CoA becomes limiting to relieve its repression on *fad* genes. Although in the RNA-Seq data, we observed a ∼1.4-fold increase in *fadR* expression in TBK-Ole, compared to TBK-Brij, this increase was CpxR-independent (Supplementary dataset S1). Using a *fadR*_200_-*lacZ* transcriptional reporter, we validated that CpxR does not regulate FadR transcriptionally; reporter expression was similar in WT and Δ*cpxR* grown in TBK-Ole (Fig 1E). These data suggested that either (i) CpxR regulates a transcriptional regulator other than FadR that must regulate *fad* genes or (ii) CpxR mediates post-transcriptional regulation of *fad* genes.

CpxR is known to regulate at least five sRNAs either directly (CyaR, CpxQ, and RprA) or indirectly (OmrA and OmrB) (34, 38, 39); any of these might have a role in Cpx-mediated post-transcriptional regulation of *fad* genes. We first investigated which of the five CpxR-regulated sRNAs are induced in LCFA-grown cells. Transcriptional reporter assays with chromosomal *lacZ* fusions of the *cis*-acting element of each sRNA as well as qRT-PCR revealed that all five sRNAs are induced in TBK-Ole-grown cells (∼2 to 6-fold) specifically in the stationary phase (Fig S6A-C). However, only the transcriptional induction of CpxQ was completely CpxR-dependent; whereas fold-induction of CyaR, OmrA, and OmrB in TBK-Ole compared to TBK-Brij was similar in WT and Δ*cpxR* strains, and RprA expression was reduced to the same extent in TBK-Brij and TBK-Ole upon *cpxR* deletion indicating that a regulator other than CpxR induces RprA in TBK-Ole, CpxQ expression was completely abolished in a Δ*cpxR* strain in both TBK-Brij and TBK-Ole (Fig S6D). Further, the CpxR-dependent increase in transcript levels was observed only for CpxQ and OmrA; whereas fold increase in transcript levels of CyaR, OmrB, and RprA in TBK-Ole compared to TBK-Brij was similar in WT and Δ*cpxR* strains, CpxQ transcript was undetected in a Δ*cpxR* strain in both TBK-Brij and TBK-Ole, and the increase in OmrA transcript levels in TBK-Ole was completely abrogated in a Δ*cpxR* strain (Fig S6E). Interestingly, the increase in transcript levels of OmrA in TBK-Ole was abolished in the Δ*cpxQ* strain, while there was no effect of *omrA* deletion on CpxQ transcript levels (Fig S6F, compare left and right panels). Collectively, of the five-known Cpx-regulated sRNAs, in LCFA-grown cells, CpxR induces the expression of only CpxQ and likely increases the stability of OmrA via CpxQ.

We next investigated whether *fad* genes are regulated by CpxQ and OmrA, the sRNAs regulated by CpxR in LCFA-grown cells (Fig S6B-F). We observed that only CpxQ deletion increased the transcript levels of *fad* genes (∼1.6 to 2.6-fold) in TBK-Ole-grown cells (Figs 2A and B). Because CyaR, OmrB, and RprA are also induced in LCFA-grown cells, although not by CpxR (Fig S6B-E), we tested whether they regulate *fad* genes as well; none of these sRNAs affected *fad* transcript levels (Fig S7A-C). The increase in *fad* transcript levels observed in the Δ*cpxQ* strain corroborated with their increase in the Δ*cpxR* strain (∼1.5 to 2.5-fold) grown in TBK-Ole (compare Figs 2A and C), and similar to the Δ*cpxR* strain, *fadE*_200_-*lacZ* transcriptional reporter was also upregulated in the Δ*cpxQ* strain (∼1.6-fold) (compare Figs 1C and 2D). Importantly, the increase in *fadE* expression in a Δ*cpxQ* strain decreased to the level observed in WT cells upon CpxQ overexpression from plasmid pZE12-*luc* (Fig 2E). CpxQ overexpression from pZE12-*luc* was validated by qRT-PCR (Fig S8). A decrease in *fadE* expression (∼25%) in WT cells upon CpxQ overexpression from pZE12-*luc* further supports the repression of *fadE* by CpxQ (Figs 2E and S8). Taken together, these data suggest that Cpx induction in the stationary phase of LCFA-grown cells represses *fad* genes via the sRNA CpxQ.

**Figure 2.**
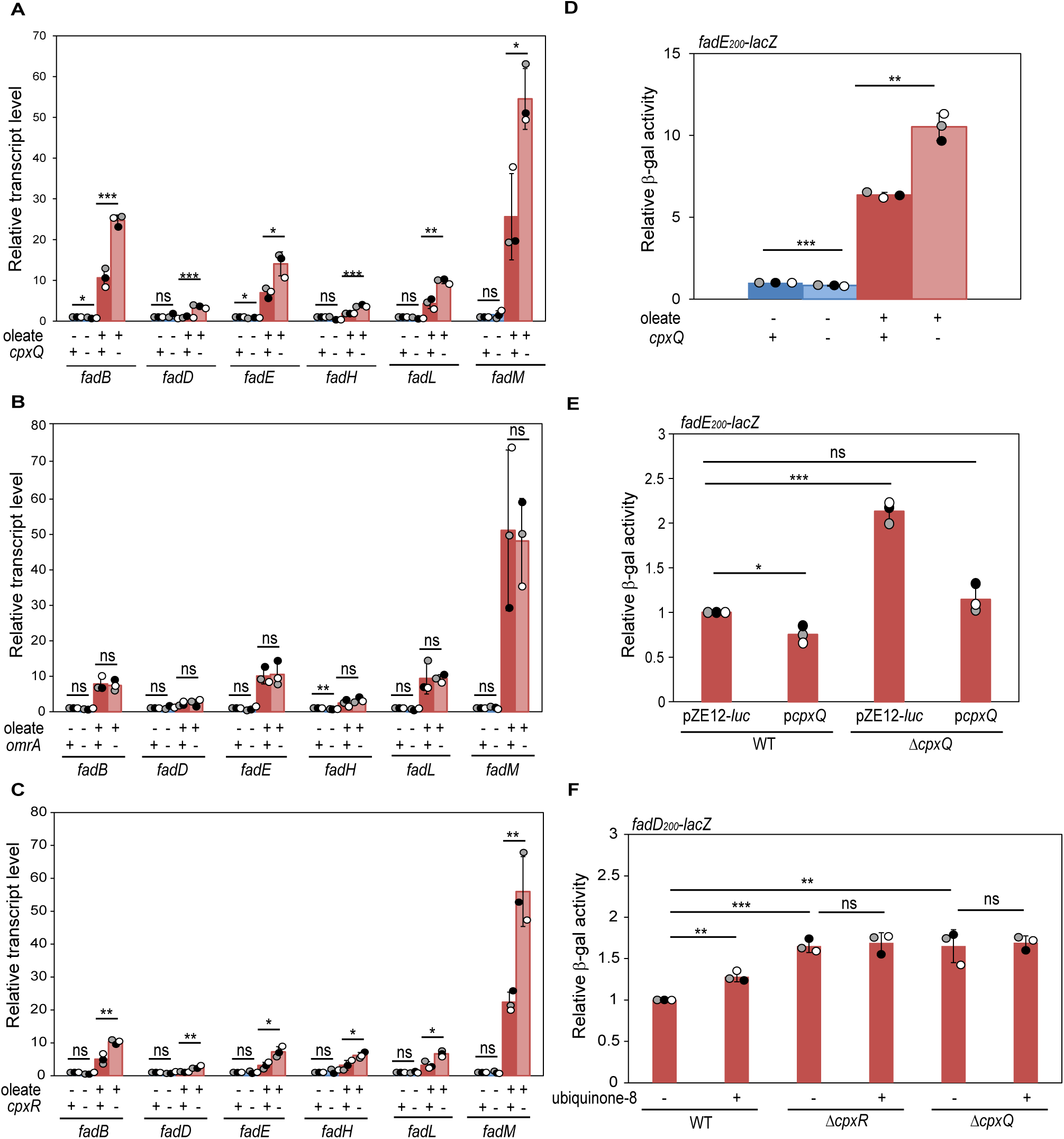
Cpx downregulates fad genes via the sRNA CpxQ. *A & B)* Amongst CpxQ (A) and OmrA (B), in oleate-grown cells, transcript levels of *fad* genes are repressed only by CpxQ. Strains were grown either in TBK-Brij or TBK-Ole till the stationary phase. Cultures were harvested for RNA isolation, cDNA was prepared, and transcript abundance was measured by qRT-PCR. Data were normalized to the transcript levels in WT in TBK-Brij and represent the average (± SD) of three independent experiments. *C)* CpxR downregulates the transcript levels of *fad* genes in oleate-grown cells. Strains were grown either in TBK-Brij or TBK-Ole till the stationary phase. Cultures were harvested and processed for qRT-PCR as mentioned in the legend to Fig 2A. Data were normalized to the transcript levels in WT in TBK-Brij and represent the average (± SD) of three independent experiments. *D)* CpxQ represses transcription of *fadE* in oleate-grown cells. Strains carrying *fadE_200_-lacZ* transcriptional reporter were grown either in TBK-Brij or TBK-Ole. Cultures were harvested in the stationary phase, and β-gal activity was measured. Data were normalized to the β-gal activity of WT in TBK-Brij. Data represent the average (± SD) of three independent experiments. The average β-gal activity (in Miller units) of WT in TBK-Brij was 96±37. *E)* The increase in *fadE* expression in a Δ*cpxQ* strain decreases to the level observed in WT cells upon CpxQ overexpression from a plasmid. Strains carrying *fadE_200_-lacZ* transcriptional reporter transformed with either pZE12-*luc* or pZE12-*luc* carrying CpxQ (p*cpxQ*) were grown in TBK-Ole. Cultures were harvested in the stationary phase, and β-gal activity was measured. Data were normalized to the β-gal activity of WT carrying pZE12-*luc*. Data represent the average (± SD) of three independent experiments. The average β-gal activity (in Miller units) of WT carrying pZE12-*luc* was 555±172. *F)* Supplementation of ubiquinone increases *fad* expression in oleate-grown WT cells. Strains carrying *fadD_200_-lacZ* transcriptional reporter were grown in TBK-Ole, supplemented with either 20 µM ubiquinone-8 or 0.1% ethanol (used for solubilizing ubiquinone-8). Cultures were harvested in the stationary phase, and β-gal activity was measured. Data were normalized to the β-gal activity of WT grown in TBK-Ole supplemented with 0.1% ethanol. Data represent the average (± SD) of three independent experiments. The average β-gal activity (in Miller units) of WT grown in TBK-Ole supplemented with 0.1% ethanol was 555±76. For panels A-F, the p-values were calculated using the unpaired two-tailed Student’s t test (***, P<0.001; **, P<0.01; *, P<0.05; ns, P>0.05).

We previously showed that ubiquinone limitation during LCFA metabolism constitutes a redox-dependent signal for Cpx activation; exogenous ubiquinone supplementation decreases Cpx activation by ∼40% in TBK-Ole-grown cells (28). To test our hypothesis that downregulation of *fad* genes by Cpx is also in response to ubiquinone limitation, we investigated whether ubiquinone supplementation decreases Cpx-mediated repression of *fad* genes. Corroborating with a ∼40% decrease in Cpx activation, exogenous supplementation of ubiquinone increased *fad* (*fadD*_200_-*lacZ*) expression by ∼30% in WT cells grown in TBK-Ole. Importantly, whereas there was a ∼1.8-fold increase in expression of *fadD* in both Δ*cpxR* and Δ*cpxQ* strains grown in TBK-Ole, there was no further effect of ubiquinone supplementation (Fig 2F). Collectively, our data suggest that the limitation of ubiquinone during LCFA metabolism is sensed by the Cpx pathway which subsequently downregulates *fad* genes via CpxQ to increase ubiquinone availability for DSB formation.

### CpxQ represses *fad* genes by downregulating FadL

Because CpxQ represses multiple *fad* genes, we considered the possibility that it mediates its effect either by increasing FadR levels post-transcriptionally or reducing its de-repression. The latter can be achieved by decreasing the acyl_(n)_-CoA pool available to bind to FadR, by decreasing LCFA transport inside the cell by downregulating FadL, and/or decreasing the conversion of LCFAs to acyl_(n)_-CoA by downregulating FadD (Fig S1). Nonetheless, in all these scenarios, CpxQ-mediated regulation will impinge on the repression of *fad* genes by FadR (Fig 1D). The increase in *fadE* expression in TBK-Ole-grown Δ*cpxR*/Δ*cpxQ*, Δ*fadR*, and Δ*cpxR*Δ*fadR/*Δ*cpxQ*Δ*fadR* strains was similar (Fig 3A and B), suggesting that CpxR/CpxQ mediates its repression on *fad* genes via FadR. Next, to address whether *fadR*, *fadD,* or *fadL* is the target of CpxQ, Fad protein levels were monitored using strains where these genes were translationally fused with a sequential peptide affinity (SPA)-tag. Whereas there was no effect of either CpxQ overexpression or deletion on protein levels of FadR and FadD in both TBK-Brij and TBK-Ole (Fig 3C and D), we observed a decrease in FadL protein levels upon CpxQ overexpression in both WT and Δ*cpxQ* strains, only in TBK-Ole (Fig 3E). These data suggest that CpxQ likely decreases LCFA transport inside the cell by downregulating FadL and as a consequence represses other *fad* genes.

**Figure 3.**
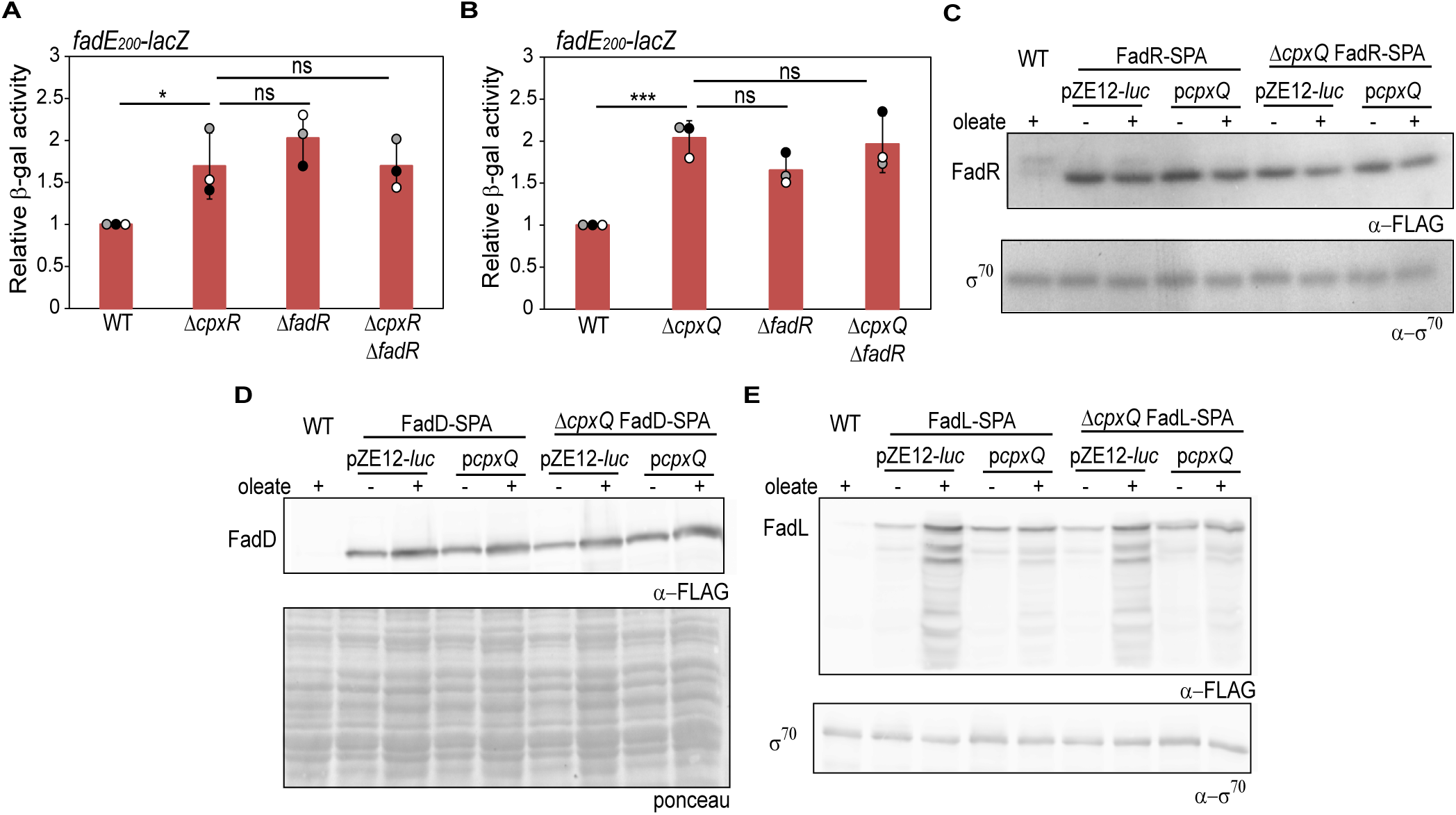
CpxQ represses fad genes by downregulating FadL. *A & B)* The repression on *fadE* exerted by CpxR (A) or CpxQ (B) is via FadR. Strains carrying *fadE_200_-lacZ* transcriptional reporter were grown in TBK-Ole. Cultures were harvested in the stationary phase, and β-gal activity was measured. Data were normalized to the β-gal activity of WT. Data represent the average (± SD) of three independent experiments. The average β-gal activity (in Miller units) of WT was 685±101 (A) and 557±56 (B). *C-E)* CpxQ overexpression decreases FadL protein levels; however it does not affect FadR and FadD levels. WT and Δ*cpxQ* strains carrying chromosomally SPA-tagged FadR (C), FadD (D), and FadL (E) transformed with either pZE12-*luc* or p*cpxQ* were grown in TBK-Brij or TBK-Ole. Cultures were harvested in the stationary phase, and processed for Western blotting. FadR-SPA (∼34 kDa), FadD-SPA (∼70 kDa), and FadL-SPA (∼57 kDa) were detected using anti-FLAG antibody. Multiple bands detected by anti-FLAG antibody in panel E (FadL-SPA) suggests degradation of FadL when overproduced. Untagged WT cells grown in TBK-Ole were used as a negative control. ο−^70^ (C & E) or ponceau S-stained counterpart of the Western blot (D) was used as a loading control. The blots are representative of three independent replicates. For panels A and B, the p-values were calculated using the unpaired two-tailed Student’s t test (***, P<0.001; **, P<0.01; *, P<0.05; ns, P>0.05).

### Besides *fad* genes, CpxQ regulates several other components of LCFA metabolism

Besides *fad* genes, our GSEA highlighted CpxR-dependent enrichment of the TCA cycle and glyoxylate shunt, gluconeogenesis, and ETC in LCFA-grown cells (Figs 1A and S4B, Supplementary datasets S1 and S2). Because CpxR regulates *fad* genes via CpxQ, we investigated whether components of other CpxR-regulated metabolic pathways are also controlled by this sRNA in LCFA condition. For this, we first validated our RNA-Seq results. Whereas CpxR deletion increased the transcript levels of candidate genes chosen from each metabolic pathway (∼1.5 to 2.5-fold) in TBK-Ole-grown cells, there was no effect in TBK-Brij (Fig 4A). Notably, amongst these candidates, only the transcript levels of *aceA* (a component of glyoxylate shunt) and *ppsA* (involved in gluconeogenesis) were ∼2.5 to 3-fold higher and those of *cyoA* and *cyoB* (components of cytochrome *bo* terminal oxidase) were ∼1.5 fold higher in Δ*cpxQ* strain in TBK-Ole (Fig 4B). We reasoned that if CpxQ represses multiple components of LCFA metabolism, then overexpression of CpxQ would compromise the growth of *E. coli* in a minimal medium containing oleate as the sole carbon source (M9+Ole). Indeed, whereas WT and Δ*cpxQ* strains could grow in M9+Ole, cells overexpressing CpxQ showed a severe growth defect (Fig 4C). Collectively, our results suggest that CpxQ decreases overall LCFA metabolism which reduces electron flow in the ETC, thereby increasing ubiquinone availability for DSB formation.

**Figure 4.**
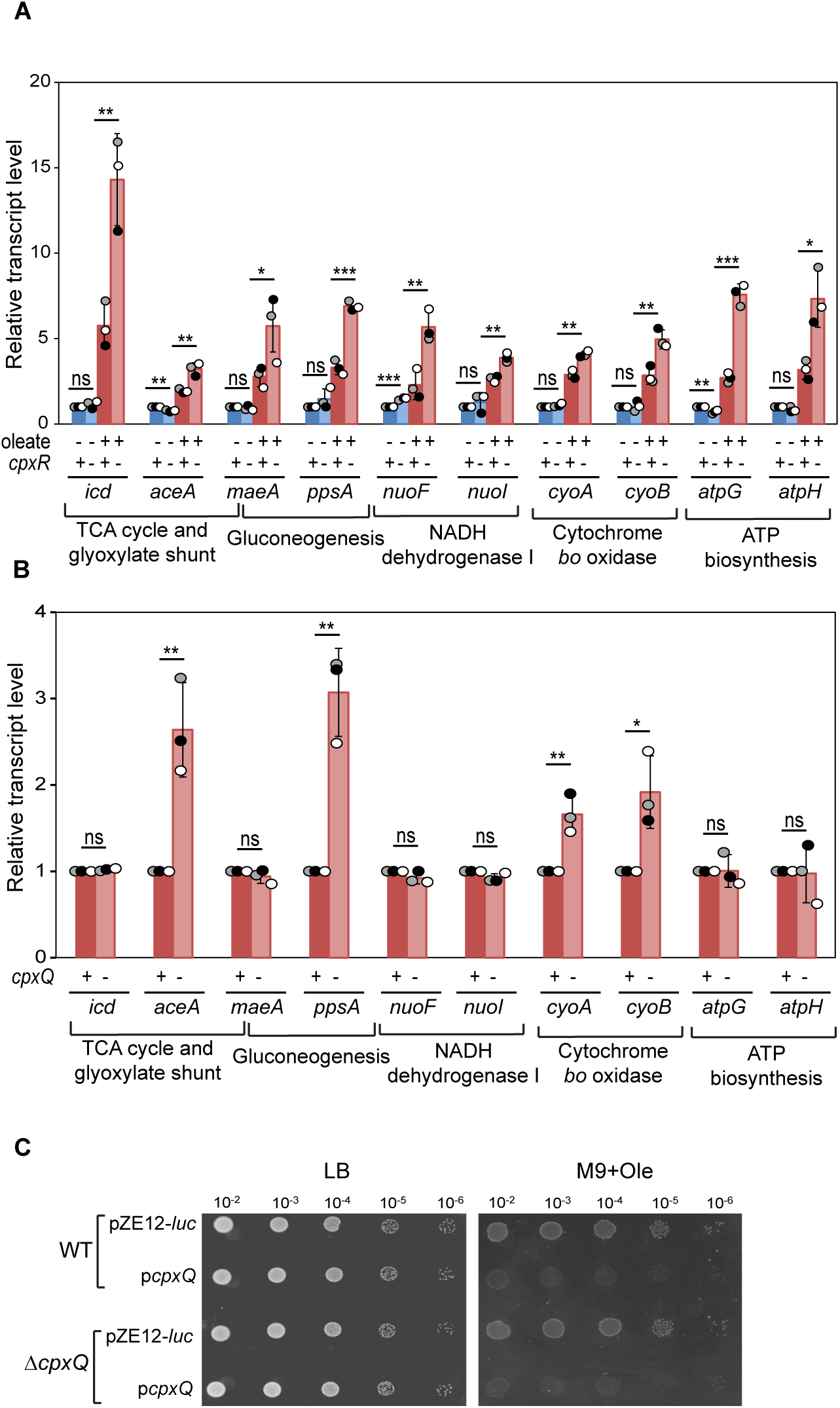
Cpx downregulates several processes involved in LCFA metabolism via CpxQ. *A)* In oleate-grown cells, CpxR downregulates components of the TCA cycle and glyoxylate shunt, gluconeogenesis, ETC, and ATP biosynthesis. Strains were grown either in TBK-Brij or TBK-Ole till the stationary phase. Cultures were harvested and processed for qRT-PCR as mentioned in the legend to Fig 2A. Data were normalized to the transcript levels in WT in TBK-Brij and represent the average (± SD) of three independent experiments. *B)* In oleate-grown cells, CpxQ downregulates a few of the CpxR-regulated components of glyoxylate shunt, gluconeogenesis, and ETC. Strains were grown in TBK-Ole till the stationary phase. Cultures were harvested and processed for qRT-PCR as mentioned in the legend to Fig 2A. Data were normalized to the transcript levels in WT and represent the average (± SD) of three independent experiments. *C)* Overexpression of CpxQ compromises the growth of *E. coli* in a minimal medium containing oleate as the sole carbon source. Strains transformed with either pZE12-*luc* or p*cpxQ* were spotted on LB and M9+Ole. For panels A and B, the p-values were calculated using the unpaired two-tailed Student’s t test (***, P<0.001; **, P<0.01; *, P<0.05; ns, P>0.05).

### Cpx increases ubiquinone levels in LCFA-grown cells via CpxQ and OmrA

Besides downregulating metabolism, another mechanism by which Cpx activation can help maintain envelope redox homeostasis in LCFA-grown cells is by increasing the oxidizing power for DSB formation by upregulating ubiquinone. Our previous work showed that total ubiquinone (ubiquinone + ubiquinol) content increases ∼1.6-fold in the stationary phase, however, this accumulation is abolished in a Δ*cpxR* strain (28). Here, we further probed the regulation of ubiquinone biosynthesis by Cpx in LCFA-grown cells. Notably, GSEA for WT grown in TBK-Brij and TBK-Ole did not show enrichment of the core ubiquinone biosynthesis pathway (comprised of proteins encoded by *ubiA* to *ubiK* and *ubiX*) that synthesizes ubiquinone from its precursors, chorismate and octaprenyl diphosphate (40, 41) (Fig S9 and Supplementary dataset S2). The differential expression analysis of various *ubi* transcripts showed that in TBK-Ole, only *ubiI* was significantly upregulated and *ubiX* was upregulated with a less significant p-value (Supplementary dataset S1 and Fig S10). Further, differential expression analysis comparing the relative transcript abundance of *ubiI* and *ubiX* in TBK-Ole-grown WT and Δ*cpxR* strains showed that their upregulation was CpxR-independent (Supplementary dataset S1). To validate these RNA-Seq results, we performed transcriptional reporter assays with strains carrying *lacZ* fused to the promoters of *ubi* genes (Fig S11A) and their transcript level analysis by qRT-PCR. In transcriptional reporter assays, whereas none of the *ubi* genes were upregulated in TBK-Ole in the exponential phase, all of them were induced in the stationary phase (Fig S11B); however, their induction was CpxR-independent (Fig S11C). Further, qRT-PCR results also corroborated the RNA-Seq data; of the twelve *ubi* genes, the transcript level of only *ubiI* and *ubiX* increased >1.5-fold in TBK-Ole in the stationary phase (compare Figs S10 and S11D); however, their increase was also CpxR-independent (Supplementary dataset S1 and S11E). Collectively, these data imply that (i) *ubi* genes are transcriptionally induced in the stationary phase by a factor other than CpxR, and (ii) CpxR increases ubiquinone levels by either upregulating genes involved in generating ubiquinone precursors (Fig S9) or regulates ubiquinone post-transcriptionally.

In *E. coli*, several metabolic pathways, including glycolysis/gluconeogenesis, pentose phosphate pathway, shikimate pathway, methylerythritol phosphate (MEP) pathway, and polyisoprenoid biosynthesis pathway contribute to the synthesis of ubiquinone precursors (40, 42) (Fig S9). Because Cpx can modulate ubiquinone levels by regulating components of the pathways that generate precursors, we searched GSEA datasets for these pathways for enrichment in TBK-Ole in a CpxR-dependent manner. GSEA for WT cultured in TBK-Brij and TBK-Ole revealed enrichment of pentose phosphate (FDR q value: 5.6%), and chorismate biosynthesis (FDR q value: 4%) pathways (Supplementary dataset S2 and Fig S3); in differential expression analysis, few components of the above pathways were upregulated in TBK-Ole (Supplementary dataset S1 and Fig S3). Amongst these pathways, GSEA for WT and Δ*cpxR* strains grown in TBK-Ole revealed CpxR-dependent enrichment of only chorismate biosynthesis (FDR q value: 6%) (Supplementary dataset S2). However, a closer look at differential expression analysis showed that none of the chorismate biosynthesis genes were upregulated in a CpxR-dependent manner; on the contrary, they were mainly <1.5-fold downregulated by CpxR (Supplementary dataset S1 and Fig S4B). Taken together, our data suggested that in LCFA-grown cells, Cpx does not regulate the transcript level of genes involved in the synthesis of either ubiquinone or its precursors.

Considering that CpxR regulates several players of LCFA metabolism via a sRNA, we next investigated the possibility of post-transcriptional regulation of ubiquinone. Because both CpxQ and OmrA, the sRNAs regulated by CpxR in LCFA-grown cells (Fig S6B-F) are dependent on Hfq, an RNA chaperone that mediates productive contacts of sRNAs with target mRNAs (39, 43), we first determined total ubiquinone content in WT, Δ*hfq,* and Δ*cpxR*Δ*hfq* strains. Whereas total ubiquinone levels were ∼1.6-fold higher in WT grown in TBK-Ole (compared to TBK-Brij), similar to the Δ*cpxR* strain, ubiquinone content decreased to basal levels in both Δ*hfq* and Δ*cpxR*Δ*hfq* strains (Fig 5A). Further, the accumulation of ubiquinone in LCFA-utilizing cells was abolished in both Δ*cpxQ* and Δ*omrA* strains (Fig 5B), which was restored when these sRNAs were expressed from a plasmid (Fig 5C and D). Similar to CpxQ, OmrA overexpression from the plasmid was also validated by qRT-PCR (Figs S8 and S12). Because CyaR, OmrB, and RprA are also Hfq-dependent sRNAs (43–45) that are induced in LCFA-grown cells, although not by CpxR (Fig S6B-E), we tested whether they regulate ubiquinone as well. None of these sRNAs affected ubiquinone levels (Fig 5B).

**Figure 5.**
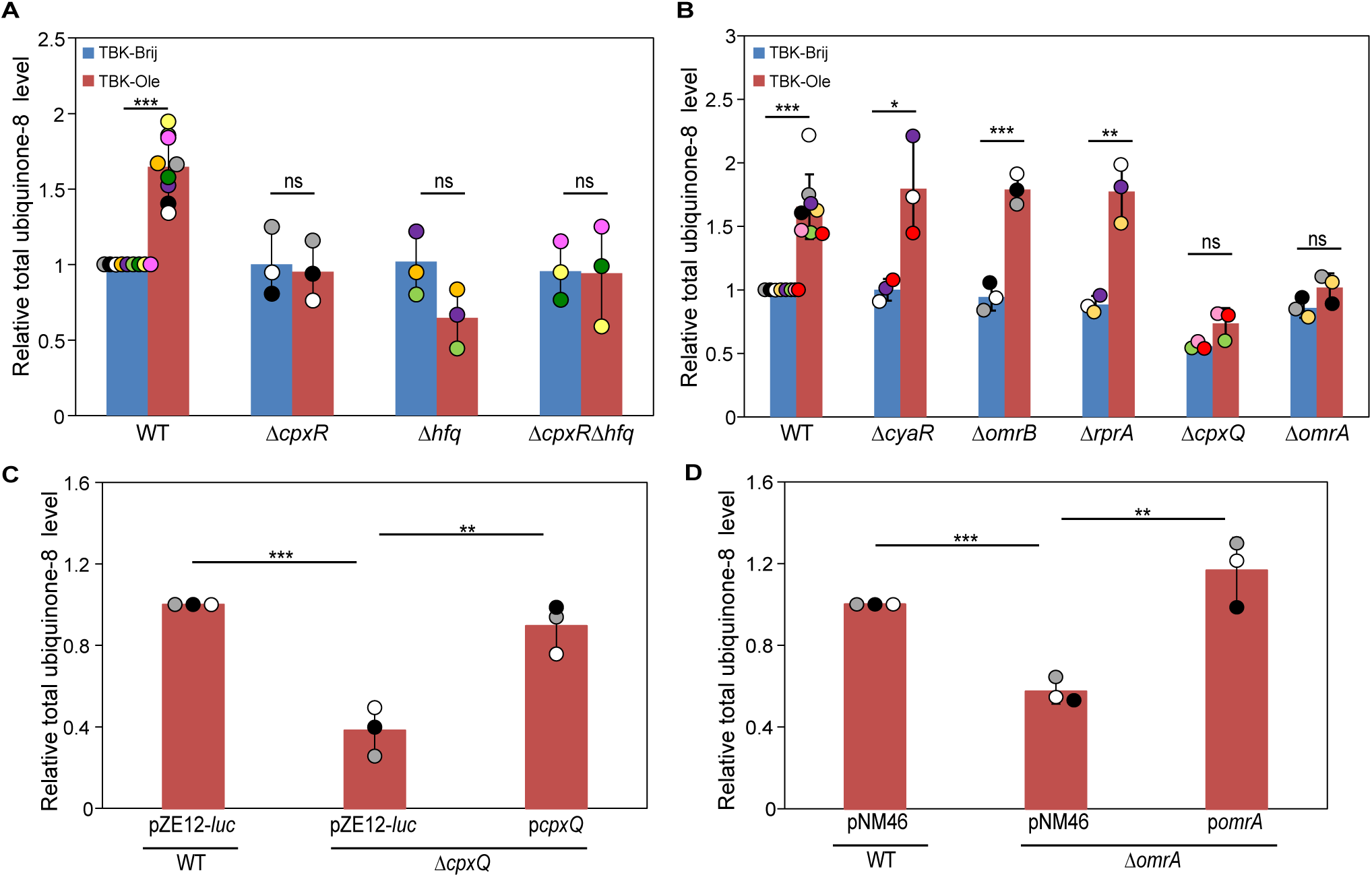
The CpxR-dependent increase in ubiquinone levels in LCFA-grown cells is mediated via CpxQ and OmrA. *A)* The increase in ubiquinone levels in oleate-grown cells is dependent on both CpxR and Hfq. Strains were grown either in TBK-Brij or TBK-Ole. The total ubiquinone-8 (ubiquinone-8 + ubiquinol-8) level in the lipid extracts was measured in the stationary phase. The total ubiquinone-8 level in each sample was normalized to the total ubiquinone-8 level of WT in TBK-Brij. Data represent the average (± SD) of at least three independent experiments. *B)* Ubiquinone levels do not increase in oleate-grown cells lacking either CpxQ or OmrA. Strains were grown either in TBK-Brij or TBK-Ole. The total ubiquinone-8 content in the lipid extracts was measured in the stationary phase. The total ubiquinone-8 level in each sample was normalized to the total ubiquinone-8 level of WT in TBK-Brij. Data represent the average (± SD) of at least three independent experiments. *C)* Ubiquinone levels are restored to WT in Δ*cpxQ* strain transformed with plasmid expressing CpxQ. Strains transformed with either pZE12-*luc* or p*cpxQ* were grown in TBK-Ole. The total ubiquinone-8 content in the lipid extracts was measured in the stationary phase. The total ubiquinone-8 level in each sample was normalized to the total ubiquinone-8 level of WT transformed with pZE12-*luc*. Data represent the average (± SD) of three independent experiments. *D)* Ubiquinone levels are restored to WT in Δ*omrA* strain transformed with plasmid expressing OmrA. Strains transformed with either pNM46 or pNM46 carrying OmrA (p*omrA*) were grown in TBK-Ole supplemented with 0.1 mM IPTG. The total ubiquinone-8 content in the lipid extracts was measured in the stationary phase. The total ubiquinone-8 level in each sample was normalized to the total ubiquinone-8 level of WT transformed with pNM46. Data represent the average (± SD) of three independent experiments. For panels A-D, the p-values were calculated using the unpaired two-tailed Student’s t test (***, P<0.001; **, P<0.01; *, P<0.05; ns, P>0.05).

In *E. coli*, the Cpx pathway is induced by a vast array of signals. Overexpression of the outer-membrane lipoprotein, NlpE, constitutes a well-known signal for Cpx activation (31, 46). Although NlpE overexpression resulted in a ∼4-fold increase in *cpxP* expression in cells grown in TBK medium, it did not increase ubiquinone levels (Fig S13A and B). Moreover, CpxQ overexpression in cells grown in TBK-Brij also did not alter ubiquinone levels (Fig S13C). These data suggest that mere activation of the Cpx pathway is not sufficient to increase ubiquinone content. We speculate that, besides Cpx activation, other LCFA-specific events occur in the stationary phase that are crucial for ubiquinone accumulation.

### CpxQ maintains redox homeostasis during LCFA metabolism

We previously demonstrated that, because LCFA-grown cells exhibit problems in DSB formation, they are sensitive to thiol agents such as DTT (28). Our present study has identified CpxQ as a major regulator of LCFA metabolism that downregulates the utilization of this carbon source and upregulates ubiquinone levels. If the CpxQ-mediated regulation on LCFA metabolism helps combat redox stress, then a Δ*cpxQ* strain should be more sensitive to DTT. Indeed, our results show that whereas WT cells can grow on TBK-Ole supplemented with 7 mM DTT and show considerable defect on 8 mM DTT, Δ*cpxQ* strain shows significant growth defect on TBK-Ole with 7 mM DTT and no growth on 8 mM DTT (Fig 6A).

**Figure 6.**
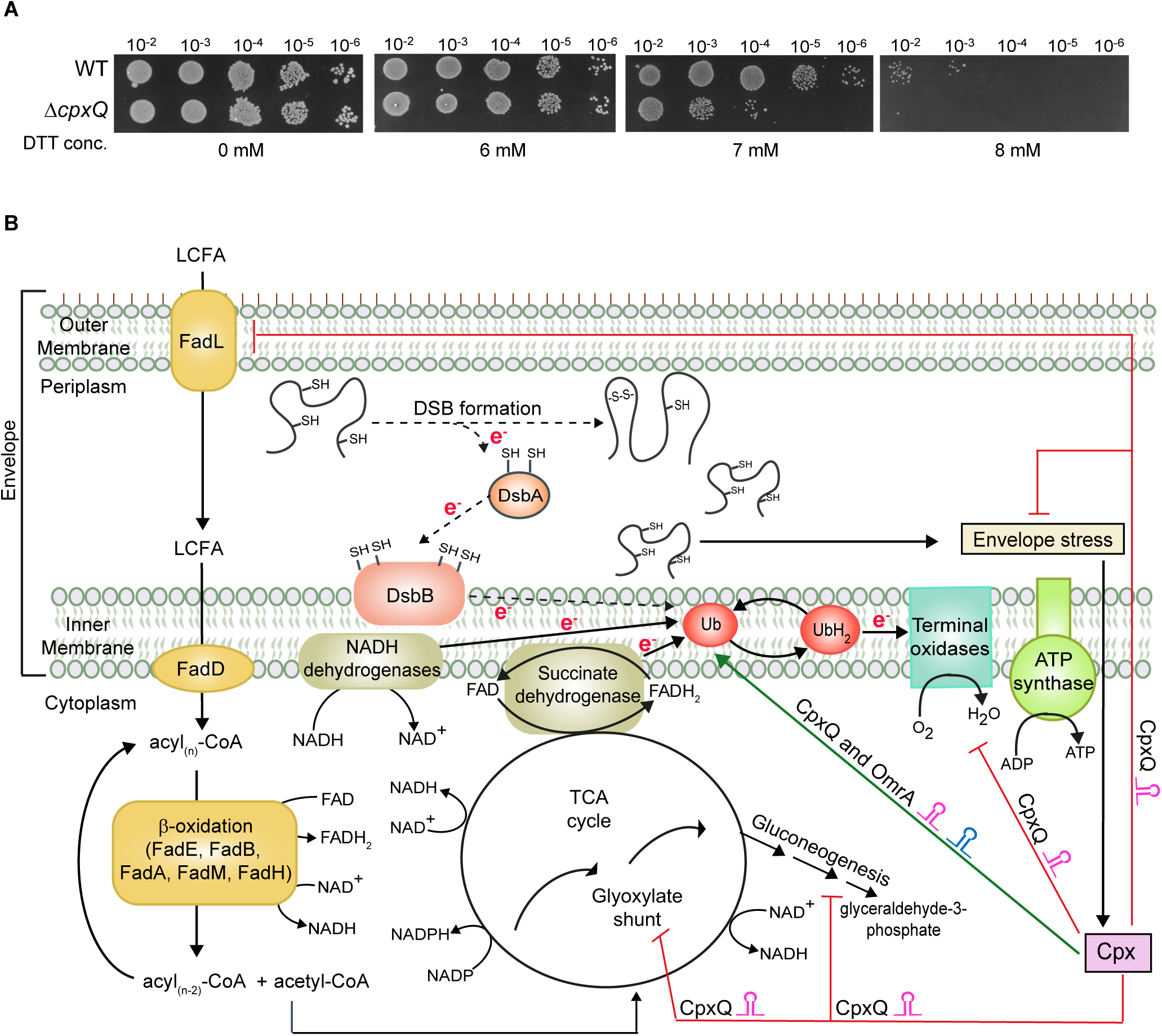
Cpx response regulates LCFA metabolism mainly via CpxQ to maintain envelope redox homeostasis. *A)* Deletion of CpxQ increases the sensitivity of *E. coli* grown in oleate to DTT. Strains were spotted on TBK-Ole with or without indicated concentrations of DTT. *B)* In LCFA-grown cells, Cpx activation facilitates DSB formation by suppressing LCFA metabolism and increasing ubiquinone content. Cpx imparts its regulation mainly via CpxQ. CpxQ downregulates various *fad* genes involved in β-oxidation, by repressing the outer-membrane LCFA transporter, FadL, and also downregulates components of glyoxylate shunt, gluconeogenesis, and ETC. CpxQ stabilizes OmrA and both sRNAs are required for ubiquinone accumulation. Arrows with *e*^-^ labeled on the line denote the direction of electron flow. The dotted arrows indicate decreased electron flow at these steps. Abbreviations: DSB, disulfide bond; LCFA, long-chain fatty acid; TCA cycle, tricarboxylic acid cycle; Ub, ubiquinone; UbH_2_, ubiquinol.

## Discussion

Despite being energy-rich, LCFAs are not the preferred nutrient source for *E. coli* likely because they confer stress in bacteria (13, 28, 47, 48). In fact, almost all pathways related to LCFA utilization are under intricate regulation by several transcriptional and post-transcriptional regulators (9–12, 14–25). Our work has identified Cpx ESR as an additional global regulator of LCFA metabolism. Prior to this study, Cpx was shown to modulate TCA cycle and ETC players in *E. coli* K-12 and EPEC (34), of which *nuo* and *cyo* operons were later confirmed as the direct targets of CpxR in EPEC (36). Our RNA-Seq in LCFA-grown *E. coli* K-12 has expanded the Cpx regulon; besides TCA cycle and ETC, in LCFA condition, glyoxylate shunt and gluconeogenesis critical for growth on non-fermentable carbon sources and LCFA transport and β-oxidation specifically required for growth in LCFAs are repressed by CpxR.

We identified several *fad* genes as the regulon members of CpxR (Figs 1C and 2C). Unlike FadR, ArcA, and cAMP-CRP, which directly control the transcription of *fad* genes (9, 10), CpxR regulates them post-transcriptionally via CpxQ (Fig 2). Our results suggest that CpxQ downregulates multiple *fad* genes by downregulating FadL; a CpxQ-dependent decrease in FadL levels (Fig 3E) would compromise the import of exogenous LCFAs, reducing acyl_(n)_-CoA pools available to bind FadR, and ultimately increasing FadR repression on *fad* genes. Our proposal is strengthened by the observation that the regulation of *fad* genes by CpxR/CpxQ is via FadR (Figs 3A and B). The control of FadR activity via CpxR/CpxQ-mediated repression of FadL likely enables FadR to simultaneously integrate information of LCFA availability and envelope stress caused by its utilization. Interestingly, FadL is repressed by another sRNA RybB, a regulon member of the ESR ο−^E^ (19). ο−^E^ is also slightly induced in LCFA-grown cells in the stationary phase (28). Because the transport of exogenous LCFAs is the first step in its metabolism, the repression of FadL by the ESRs Cpx and ο−^E^ via sRNAs CpxQ and RybB, respectively, might serve as an important checkpoint to decrease overall LCFA metabolism to counteract envelope stress.

Besides *fad* genes involved in LCFA transport and β-oxidation, we confirmed CpxR-mediated repression of several genes representing the other LCFA-related pathways (Fig 4A). CpxR possibly regulates these candidate genes by distinct mechanisms. In *E. coli* K-12, we identified a CpxR binding site upstream of the transcription start site of *nuoA* (the first gene of the *nuo* operon) and *cyoA* (the first gene of the *cyo* operon) (Fig S14). Thus, similar to EPEC, CpxR likely directly represses *nuo* and *cyo* operons in *E. coli* K-12 as well. However, our results that *cyoA* and *cyoB* are also repressed by CpxQ (Fig 4B), suggest that during LCFA metabolism, CpxR-CpxQ-*cyo* forms a feed-forward loop that enables tight repression of the terminal oxidase. Besides *cyoA* and *cyoB*, CpxQ downregulates *aceA* and *ppsA* (Fig 4B). Because *iclR*, the repressor of the *aceBAK* operon (49), is transcriptionally activated by apo-FadR, we speculate that CpxQ represses *aceA* via FadR. The other four genes (*icd*, *maeA*, *atpG,* and *atpH*) repressed by CpxR (Fig 4A), are neither regulated by CpxQ (Fig 4B), nor show any putative CpxR binding site within ∼500 bp upstream of the translational start site, indicating that they are regulated by CpxR via another intermediary factor.

Besides downregulating LCFA degradation, Cpx increases ubiquinone levels via the sRNAs, CpxQ and OmrA (Fig 5). In *E. coli*, several *ubi* genes are regulated by transcriptional regulators mainly ArcA, FNR, and cAMP-CRP, under growth conditions, such as oxygen and carbon source availability (15, 16, 50–53). However, there are conflicting reports on the regulation of ubiquinone by sRNAs. The 3’ end of *ubiJ* locus was proposed to encode a sRNA EsrE (54); whereas one study reported the involvement of both UbiJ protein and the sRNA EsrE in ubiquinone biosynthesis (55), other studies showed that only UbiJ protein is involved in this process (56, 57). Very recently, a sRNA SicX, strongly induced by low-oxygen conditions has been shown to positively regulate ubiquinone in *P. aeruginosa* by activating the translation of *ubiUVT* mRNA responsible for anaerobic ubiquinone biosynthesis (58). Because aerobic ubiquinone biosynthesis in *E. coli* requires several metabolic processes for the generation of precursors followed by a dedicated pathway comprised of at least twelve *ubi* genes (Fig S9), multiple steps might be controlled by CpxQ and OmrA to increase ubiquinone content in LCFA-grown cells. Based on the observations that CpxR does not increase the transcript levels of *ubi* genes or genes involved in generating ubiquinone precursors (Figs S11E and Supplementary datasets S1 and S2) but Cpx-regulated sRNAs positively regulate ubiquinone (Fig 5B-D), we speculate that CpxQ and OmrA increase the translation of players of ubiquinone biosynthesis. Because all *ubi* genes are transcriptionally induced in LCFA-grown cells (Fig S11B) but there is no increase in their transcript levels (except *ubiI* and *ubiX*) (Figs S10 and S11D), sRNAs are likely required for the rapid translation of the newly synthesized transcripts. Given the complexity of the ubiquinone biosynthesis pathway, a combination of high-throughput approaches enabling the co-purification of target mRNAs with sRNAs and comparative proteome analysis of cells carrying or lacking sRNAs will be required to identify the direct/indirect targets of CpxQ and OmrA (59). Because Cpx activation alone does not increase ubiquinone levels (Fig S13), these high-throughput techniques must be performed in the presence of LCFAs. In fact, pentose phosphate and chorismate biosynthesis pathways, that provide ubiquinone precursors, are upregulated in LCFA-grown cells (Fig S3 and Supplementary datasets S1 and S2). Upregulation of these pathways, although in a CpxR-independent manner (Fig S4B and Supplementary datasets S1 and S2), might be crucial for CpxQ and OmrA to act at the downstream steps.

Cpx response impacts every aspect of the LCFA metabolic pathway to facilitate DSB formation by increasing ubiquinone availability (Fig 6B). Contrary to its conventional mode of imparting regulation via CpxR working as a transcriptional regulator, in LCFA-grown cells, Cpx regulates the metabolic players mainly post-transcriptionally via the sRNA CpxQ. The vital role of CpxQ in maintaining envelope redox homeostasis is evident from the following observations: i) exogenous supplementation of ubiquinone decreases CpxQ-mediated repression of *fad* genes (Fig 2F), suggesting that CpxQ increases ubiquinone availability by suppressing LCFA metabolism, ii) CpxQ stabilizes OmrA and both sRNAs directly increase ubiquinone content (Figs S6F and 5B-D), and iii) *cpxQ* deletion sensitizes LCFA-grown *E. coli* to thiol stress (Fig 6A). The preventive mode of Cpx in maintaining envelope redox balance by modulating metabolism in LCFA-grown cells is in contrast to its remedial mode of restoring envelope integrity by upregulating protein quality control factors observed in other Cpx activating conditions (32, 34). Except *degP*, transcript levels of none of the other known Cpx-dependent envelope-localized chaperones and proteases (*cpxP*, *dsbA*, *spy*, *ppiA*) increase in LCFA-grown cells (Supplementary Dataset S1).

*M. tuberculosis*, *P. aeruginosa*, *S.* Typhimurium, and *V. cholerae* use host-derived LCFAs, which contributes to their survival and virulence (1–8). Notably, DSB formation, crucial for the functioning of several virulence factors, occurs in the envelope of these pathogens (26, 60–63). Therefore, it is imperative to ascertain whether these bacteria are also predisposed to LCFA-induced redox stress and, if so, whether they use defense mechanisms similar to *E. coli*. Interestingly, in *V. cholerae*, LCFAs induce the expression of a sRNA FarS that reinforces repression on two *fadE* paralogs by FadR (64). Further, the transcriptome of *M. tuberculosis* cultured in LCFAs showed increased expression of genes related to reductive stress as well as high expression from the intergenic regions, suggesting a link between redox stress and sRNA-mediated control (65). It would be exciting to investigate whether sRNAs play a role in governing envelope redox balance during LCFA metabolism in these pathogens. The incorporation of sRNAs as an integral part of the LCFA-induced ESRs would provide a fast response to the stimulus and quick recovery after the removal of the stimulus.

## Materials and Methods

### Strains, plasmids, and primers

The strains and plasmids, and primers used in this study are listed in Supplemental Tables S3 and S4, respectively. *E. coli* BW25113 was used as the WT strain for various assays. Deletion strains obtained from either Keio collection (66) or ASKA gene knock-out library (67) were freshly transduced using P1 phage and verified by colony PCR before final assays, to rule out genetic errors. Wherever required, the antibiotic resistance cassette from deletion strains was flipped out using pCP20. The derivatives of *cpxR*, *cpxQ*, and *hfq* deletions were unstable as glycerol stocks and therefore, these deletions were always freshly transduced. Strain DH5α was used for cloning in pZE12-*luc* and pNM46, and BW25142 (*pir*+) was used for cloning in pAH125.

The chromosomal single-copy transcriptional reporter fusions were constructed using the pAH125 integration vector, as described previously (28, 68). Briefly, the *cis*-acting element of the gene was PCR amplified from the genomic DNA of BW25113 and cloned upstream of *lacZ* at *Kpn*I/*Sal*I and *Eco*RI sites of pAH125. For each reporter fusion, except *cpxP*-*lacZ*, cloned DNA fragments comprised of ∼200-1000 bp upstream and ∼18-50 bp downstream of the translational start site. For *cpxP*-*lacZ*, the cloned DNA fragment comprised of 410 bp upstream and 223 bp downstream of the translational start site. For all reporter fusions, the approximate length of the DNA fragment cloned in pAH125 is denoted by a subscript (after the gene name) in the relevant genotype column in Table S3. The cloned plasmid was further integrated at the *att*λ site of BW25113 using integrase expressed from the pINT-ts helper plasmid. Single copy integrants were PCR verified. The integrants were P1 transduced into clean BW25113 before use.

The sRNA deletion and SPA-tagged strains were constructed using the λ Red-recombinase system (69, 70), using a helper plasmid (pSH06) expressing λ Red-recombinase enzymes. pSH06 was modified from pSim5 plasmid where the chloramphenicol resistance gene was replaced by ampicillin resistance gene (amplified from pKD46) at the *Eco*RI and *Nco*I sites. Briefly, for deletion strains, primers that carried sequences homologous to the sites adjacent to the sRNA to be deleted were used to amplify antibiotic resistance cassette, flanked by flippase recognition target (FRT), either from pKD3 (harbors chloramphenicol resistance cassette) or pKD13 (carries kanamycin resistance cassette). For tagging the C-terminus end of the gene at the chromosomal locus with SPA, the sequence of SPA along with the kanamycin cassette was PCR amplified from a SPA-tagged strain obtained from the Keio collection (66) using primers that carried sequences homologous to the target gene. The amplified fragment was transformed into BW25113 expressing λ Red-recombinase enzymes from pSH06. The recombinants were selected using the desired antibiotic and PCR verified. The deletions or the SPA-tagged loci were P1 transduced into clean BW25113 before use.

For cloning CpxQ and OmrA in plasmids, these sRNAs were amplified from BW25113 genomic DNA. CpxQ was cloned at the *Xba*I site downstream of luciferase in pZE12-*luc* to yield pMK05. OmrA was cloned at the *Aat*II and *Eco*RI sites in pNM46 to yield pNV13.

### Media composition and growth conditions

The media used had the following composition: lysogeny broth (LB), 5 g/liter Bacto yeast extract, 10 g/liter Bacto tryptone, and 5 g/liter NaCl; tryptone broth K (TBK), 10 g/liter Bacto tryptone, and 5 g/liter KCl; M9 minimal medium, 5.3 g/liter Na_2_HPO_4_, 3 g/liter KH_2_PO_4_, 0.5 g/liter NaCl, 1 g/liter NH_4_Cl, 0.12 g/liter MgSO_4_, 2 mg/liter biotin, 2 mg/liter nicotinamide, 0.2 mg/liter riboflavin, and 2 mg/liter thiamine. TBK medium was buffered at pH 7.0 using 100 mM potassium phosphate. Where required, medium was supplemented with sodium oleate as a carbon-source, at the final concentration of 5 mM. 50 mM stock of oleate was prepared in 5% Brij-58, filter-sterilized and stored at -20°C. Sodium oleate and Brij-58 were obtained from Sigma. 1.5% (w/v) Difco agar was used for media solidification. When required, antibiotics, ampicillin (100 μg/ml), chloramphenicol (20 μg/ml), kanamycin (30 μg/ml), and spectinomycin (50 μg/ml), were used.

Primary cultures were set up in 3 ml LB. These were used to further set up secondary cultures in 15 ml TBK medium supplemented with either 0.5% Brij-58 (TBK-Brij) or 5 mM oleate (TBK-Ole) in a 125 ml flask with an initial OD_600_ of ∼0.01. When required, the medium was supplemented with 20 µM ubiquinone-8 (Avanti Polar Lipids). 20 mM stock of ubiquinone-8 was prepared in 100% ethanol (Merck). Cultures were grown in a water bath shaker at 37°C and 220 rpm.

### Growth curves

Growth curves were carried out in shake flasks. Primary cultures in LB were pelleted, washed, and resuspended in TBK medium. Secondary cultures were set up in TBK-Brij or TBK-Ole by re-inoculating cells to an initial OD_600_ ∼0.01. Cultures were grown and OD_600_ was measured at required time intervals.

### β-galactosidase assay

β-gal assays were performed in either exponential or stationary phase, as described previously (28). Briefly, cells were pelleted, washed thrice with Z-buffer, and re-suspended in the same buffer to an OD_450_ ∼0.5. Promoter activity was measured by monitoring β-gal expression from chromosomal transcriptional reporter fusions, as mentioned (71).

### RNA isolation, cDNA preparation, and quantitative RT-PCR

Secondary cultures were harvested at either the exponential or stationary phase. RNA was isolated by the hot-phenol method and cDNA was prepared, as described previously (13). Briefly, 8 ml cultures were added to ice-cold 5% water-saturated phenol (in ethanol), and centrifuged at 8200 X g for 2 min. Cell pellets were either flash-frozen in liquid nitrogen and stored at -80°C until required, or directly processed for RNA extraction. Cell pellets were re-suspended in 500 µl of lysis solution (320 mM sodium acetate pH 4.6, 8% SDS, 16 mM EDTA), and mixed with 1 ml water-buffered phenol. Samples were heated at 65°C for 5 min with intermittent vortexing after every 40 sec. Samples were kept on ice for 5 min and centrifuged at 14000 X g at 4°C for 10 min. The supernatant was extracted twice with phenol-chloroform, followed by precipitation with 2.5 volumes of absolute ethanol at 4°C. The RNA pellet was washed with 70% ethanol, air-dried, and resuspended in 85 µl nuclease-free water. Genomic DNA was removed by DNA-free Turbo DNase according to the manufacturer’s instructions for rigorous DNase treatment (Applied Biosystems, USA). 5 µg RNA was used to prepare cDNA for qRT-PCR using SuperScript III reverse transcriptase (Applied Biosystems, USA).

The qRT-PCR reactions were set up using Power Sybrgreen PCR master mix following the manufacturer’s instructions (Applied Biosystems, USA), and 5 pmol of forward and reverse primers. Real-time PCR was performed with either Quant Studio 6 Flex system (Applied Biosystems, USA) or MasterCycler EP Realplex 4 thermal cycler (Eppendorf). Data were analyzed using *recA* and *gyrA* as internal controls, as described (72).

### RNA sequencing and initial bioinformatic analysis

RNA sequencing was performed on four distinct samples, WT BW25113 and its isogenic Δ*cpxR* strain, in both TBK-Brij and TBK-Ole (WT_TBK-Brij, WT_TBK-Ole, Δ*cpxR*_TBK-Brij, and Δ*cpxR*_TBK-Ole), cultured to stationary phase. For each sample, three biological replicates were processed. RNA was extracted as described above for qRT-PCR. To ensure purity, before genomic DNA removal, isolated RNA was passed through a column from the RNeasy Micro kit (Qiagen), following the manufacturer’s instructions. The quality and integrity of RNA were assessed by RNase-free gel electrophoresis. RNA sequencing was outsourced to MedGenome Labs Ltd., India. The company performed the quality control of the RNA samples and ribosomal RNA depletion followed by cDNA library preparation. Pair-end sequencing was performed using prepared cDNA libraries on Illumina HiSeqX to generate 60 million reads, 2×150bp reads/sample. The sequenced data comprised twelve RNA sequencing read files in fastq.gz format.

Quality assessment of the sequencing data was performed using MultiQC (73). Subsequently, the reads were aligned to the reference genome using the STAR aligner (version 2.7.10b), employing the genome sequence of *E. coli* BW25113 (RefSeq ID: NZ_CP009273.1). The resultant alignment files were in .bam format (74). Quantification of features was conducted using the Subread package (75), which tallies the reads mapping to specific genes. Following feature counting, raw counts were normalized, and differential expression analysis was performed employing the DESeq2 (76) package in the R environment. The input files for this analysis included the raw read counts derived from the .bam files processed through Subread, as well as a metadata file categorizing the samples according to their respective types (WT or Δ*cpxR*) and growth conditions (TBK-Brij or TBK-Ole). The output from DESeq2 was utilized to generate volcano plots using R scripts for the following comparisons: WT_TBK-Ole vs. WT_TBK-Brij, Δ*cpxR*_TBK-Brij vs. WT_TBK-Brij, and Δ*cpxR*_TBK-Ole vs. WT_TBK-Ole.

### Gene set enrichment analysis and generation of heat map

GSEA was performed using version 4.3.2 of the GSEA tool (77). Input data for GSEA comprised read counts (transcripts per million), phenotype labels, and a gene set file derived from the EcoCyc database of *E. coli* K-12 MG1655, which contained a total of 399 gene sets (78). Additionally, two supplementary pathways, namely the methylerythritol phosphate pathway I and LCFA transport and β-oxidation, were appended, resulting in a total of 401 pathways. Gene sets with a minimum of three genes were considered for analysis. The comparison groups for GSEA were: WT_TBK-Ole vs. WT_TBK-Brij, Δ*cpxR*_TBK-Brij vs. WT_TBK-Brij, and Δ*cpxR*_TBK-Ole vs. WT_TBK-Ole.

A heat map representing the difference in gene expression of pathways in the four sample types (WT_TBK-Brij, Δ*cpxR*_TBK-Brij, WT_TBK-Ole, and Δ*cpxR*_TBK-Ole) was generated from normalized data obtained from DESeq2 using HemI 1.0 (Heatmap Illustrator) tool (79).

### Western blotting

The expression of SPA-tagged Fad proteins was monitored by Western blot analysis. Cultures were grown to stationary phase and an equal number of cells (OD_600_ ∼1) were pelleted, resuspended in protein loading buffer, and incubated at 95°C for 10 min for lysis. Cell lysates were electrophoresed on SDS-PAGE and proteins were transferred to a nitrocellulose membrane. Equal loading of samples was monitored by staining the membrane with PonceauS. An image of the Ponceau-stained membrane was captured by the GelDoc Go Imaging System (Bio-Rad). The membrane was blocked with 5% (wt/vol) skim milk at 4°C for 12-16 hours and probed with either anti-FLAG (1:3300; Sigma) or anti-ο−^70^ (1:3300; BioLegend) primary antibody and horseradish peroxidase-conjugated anti-mouse (1:5000; Sigma) secondary antibody. Chemiluminescence signals were developed with SuperSignal West Dura extended-duration substrate (Pierce) and were captured with either ImageQuant LAS4000 imager (GE Healthcare) or on an X-ray film.

### Dilution spotting

Primary cultures were pelleted, washed, and re-suspended in M9 minimal medium without any carbon source and normalized such that all samples had the same number of cells. Different dilutions of the cultures were spotted on the desired solid media. Plates were incubated and imaged at various time intervals using GelDoc Go Imaging System (Bio-Rad).

### Quinone extraction and detection by HPLC-photodiode array detection analysis

Quinones were extracted as described previously (13). An equal number of cells (∼3 × 10^10^) were pelleted and pellet mass was determined. Pellet was re-suspended in 100 μl 0.15 M KCl. 200 μl glass beads (acid washed ≤ 106 µm, Sigma), 600 μl methanol, and 12 μg ubiquinone-10 standard (Sigma) in hexane (used as control for extraction efficiency) were added to the re-suspension. Samples were vortexed for 15 min followed by the addition of 400 μl hexane. Samples were again vortexed for 3 min followed by centrifugation at 6000 rpm for 1 min. The upper hexane layer (∼200 μl) was transferred to a fresh microcentrifuge tube, completely dried under vacuum, and re-suspended in 100 μl mobile phase (isocratic solution comprised of 40% ethanol, 40% acetonitrile, and 20% mixture of 90% isopropyl alcohol and 10% 1 M lithium perchlorate). Samples were separated by reverse-phase HPLC using a C18 column (Waters Sunfire 5 μm column, 4.6 × 250 mm). The samples were injected at a flow rate of 1 ml/min along with the mobile phase at a temperature of 25°C. Quinones were detected using a Photodiode array detector. The peaks for ubiquinone-8 (λ_max_ 275 nm) and ubiquinol-8 (λ_max_ 290 nm) in the effluent were assigned based on the elution time of pure standards. Ubiquinol-8 was prepared by the reduction of ubiquinone-8 following the procedure described in (13). The total ubiquinone content in the sample was calculated by the addition of the peak area per unit mass for ubiquinone-8 and ubiquinol-8, which was divided by the ubiquinone-10 peak area to account for the difference in extraction efficiency between the samples.

## Data Availability

The raw RNA-sequencing data are available at the NCBI’s SRA database. The remaining data are available in the main article and its supporting information.

## Acknowledgments

We acknowledge Christopher Rao, Don Court, Gisela Storz, Jean-François Collet, and Kai Papenfort for strains and plasmids. We thank Apuratha Pandiyan for assistance with the initial bioinformatics analysis of RNA-Seq data. We acknowledge Navjot Kaur for constructing plasmid pNV13 and Saloni Sahu for making strains RC25028 and RC25029. This work was supported by the Department of Biotechnology (DBT)/Wellcome Trust India Alliance senior fellowship/grant IA/S/21/2/505907 and partially supported by Ministry of Education-Scheme for Transformational and Advanced Research in Sciences (STARS) grant MoE-STARS/STARS-1/296, Department of Science and Technology-Science and Engineering Research Board (DST-SERB) grant CRG/2018/000833, DBT grant BT/PR34553/BRB/10/1846/2020 and funds from IISER Mohali (to R.C.), and DST-INSPIRE Faculty Award and Seed Grant from IIT Hyderabad (to G.S.). M.S. and D.R. acknowledge IISER Mohali, M.K. thanks DBT, S.S. acknowledges Council of Scientific and Industrial Research (CSIR), and V.S. and R.A.K thank University Grants Commission (UGC) for fellowship support for doctoral work.

