## Supporting Information for "An envelope stress response governs long-chain fatty acid metabolism via a small RNA to maintain redox homeostasis in *Escherichia coli*"

**This PDF file includes:**

Figures S1 to S14
Tables S1 to S4
Legends for Datasets S1 and S2
SI References

**Other supporting materials for this manuscript include the following:**

Datasets S1 and S2

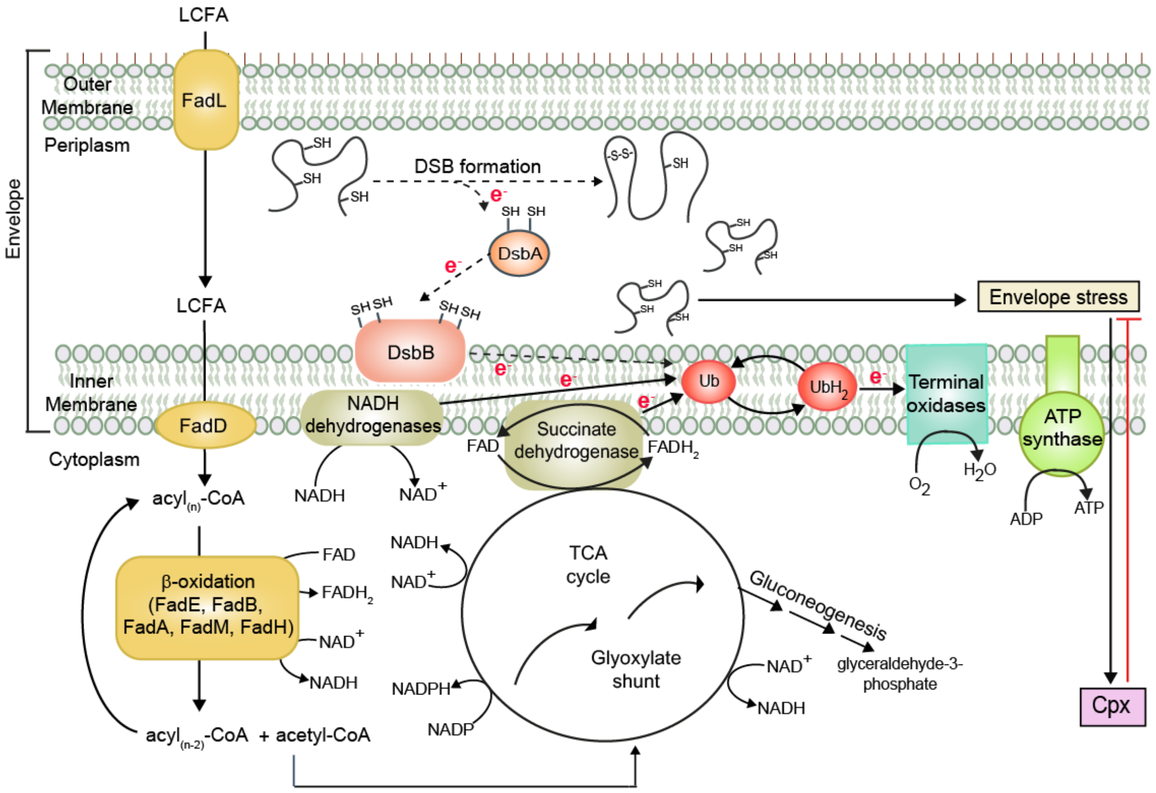

**Fig. S1.** *LCFA metabolism causes envelope stress and induces Cpx response as a combat strategy.* Exogenous LCFAs are transported across the outer membrane by FadL and further extracted from the inner membrane by the acyl-CoA synthetase, FadD, which also esterifies them to acyl$_{(n)}$-CoA. Acyl$_{(n)}$-CoAs are degraded to acetyl-CoA via β-oxidation mediated by the activity of several Fad enzymes. Acetyl-CoA feeds into central metabolism, and NADH and $FADH_2$ produced during metabolism are oxidized in the ETC for energy generation (please see text for details). LCFA metabolism increases electron flow towards ubiquinone in the ETC, limiting ubiquinone for DSB formation in secreted proteins (performed by the DsbA-DsbB machinery), thereby causing envelope stress. *E. coli* induces the Cpx response, which restores homeostasis. Arrows with $e^-$ labeled on the line denote the direction of electron flow. The dotted arrows indicate decreased electron flow at these steps. Abbreviations: DSB, disulfide bond; LCFA, long-chain fatty acid; TCA cycle, tricarboxylic acid cycle; Ub, ubiquinone; $UbH_2$, ubiquinol.

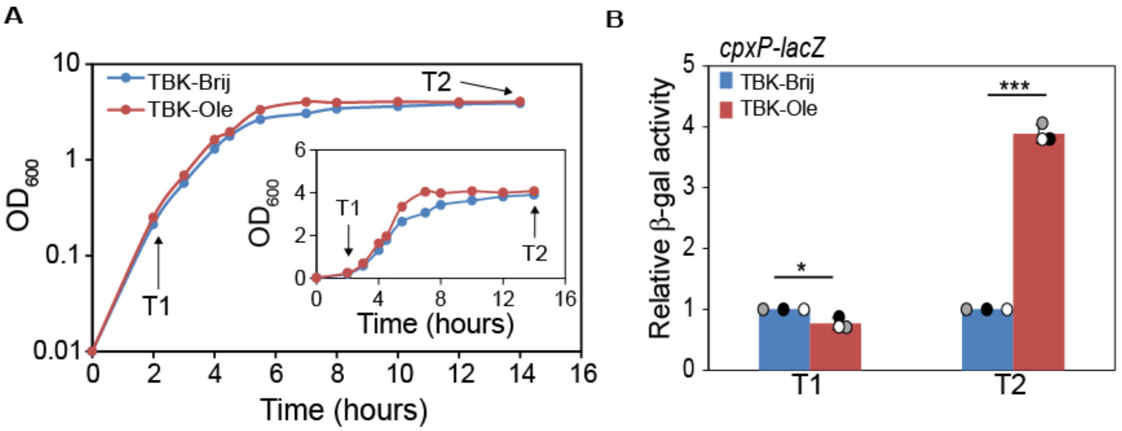

**Fig. S2.** *Cpx is induced in LCFA-grown E. coli BW25113 in the stationary phase. A)* Growth curve of WT in TBK-Brij and TBK-Ole. $OD_{600}$ of the cultures was measured. Growth curves were plotted on a semi-logarithmic scale. T1 and T2 denote exponential and stationary phase time points, respectively, where cultures were harvested for various assays. Inset: The growth curves were also plotted on a linear scale. *B)* Cpx is induced in the stationary phase in oleate-utilizing cells. WT carrying *cpxP-lacZ* transcriptional reporter was grown either in TBK-Brij or TBK-Ole. Cultures were harvested in the exponential and stationary phases of growth. β-gal activity was measured, and data were normalized to the β-gal activity of WT grown in TBK-Brij at respective phases. Data represent the average (± SD) of three independent experiments. The average β-gal activity (in Miller units) of WT in TBK-Brij at exponential and stationary phases was 644±143 and 169±11, respectively. For panel B, the p-values were calculated using the unpaired two-tailed Student's t test (***, P<0.001; **, P<0.01; *, P<0.05; ns, P>0.05).

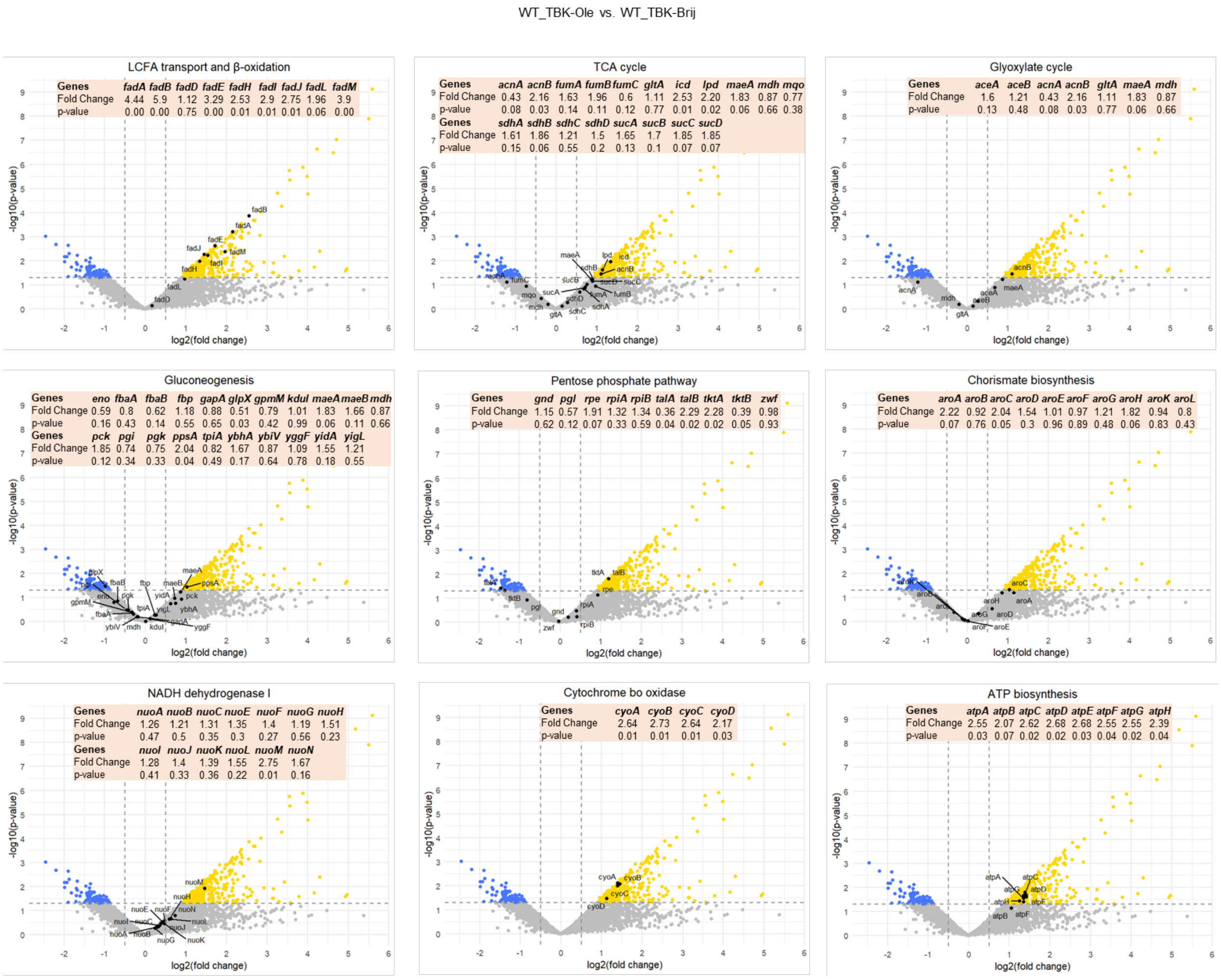

**Fig. S3.** *Volcano plots for the LCFA-related pathways enriched in the GSEA dataset for the comparison group WT_TBK-Ole vs WT_TBK-Brij*. The significant range of log2 fold change values was -0.5 to 0.5 and the p-value cut-off was 0.05. The golden yellow dots represent upregulated genes, blue dots represent downregulated genes, and the grey dots represent genes with insignificant expression given the threshold values considered for generating the plots. Black dots indicate genes of the respective pathways as mentioned on the top of each volcano plot. The table on the top represents the fold change values and p-values for all genes in the respective pathway.

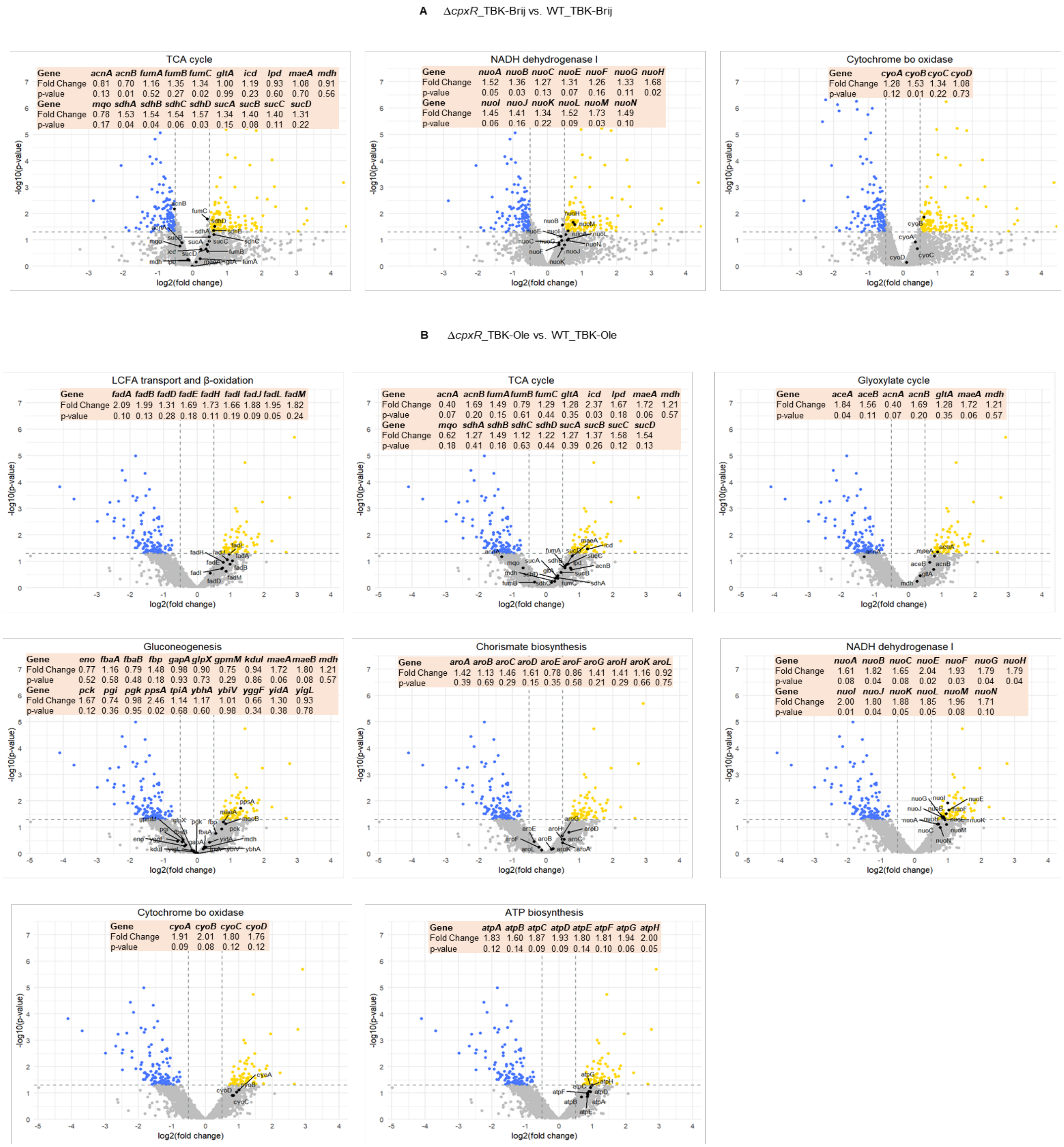

**Fig. S4.** *Volcano plots for the LCFA-related pathways enriched in the GSEA dataset for the comparison group ΔcpxR_TBK-Brij vs WT_TBK-Brij (A) and ΔcpxR_TBK-Ole vs WT_TBK-Ole (B), respectively.* The significant range of log2 fold change values was -0.5 to 0.5 and the p-value cut-off was 0.05. The golden yellow dots represent upregulated genes, blue dots represent downregulated genes, and the grey dots represent genes with insignificant expression given the threshold values considered for generating the plots. Black dots indicate genes of the respective pathways as mentioned on the top of each volcano plot. The table on the top represents the fold change values and p-values for all genes in the respective pathway.

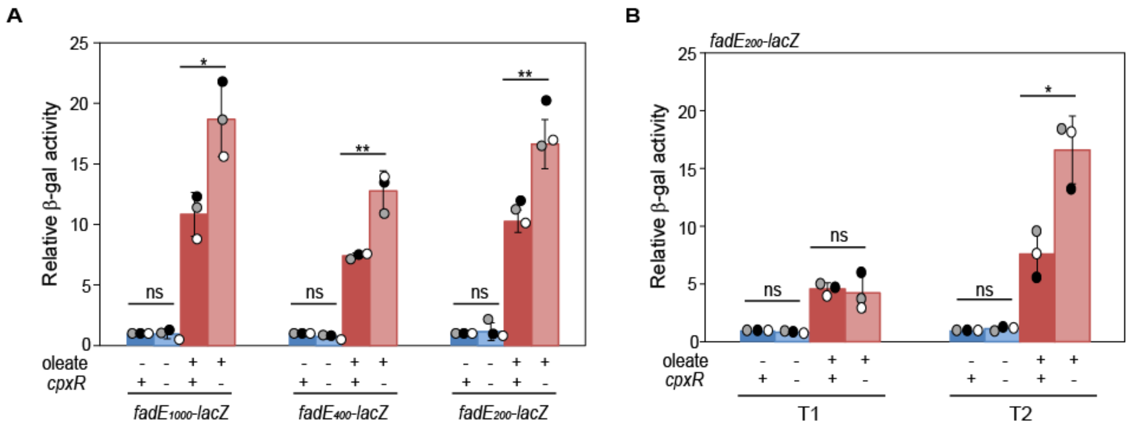

**Fig. S5.** *The site required for regulation of fadE by Cpx lies in its ~200 bp cis*-acting *element. A)* Truncation analysis reveals that ~200 bp region upstream of *fadE* harbors the site regulated by Cpx. Strains carrying *fadE1000-lacZ*, *fadE400-lacZ*, and *fadE200-lacZ* transcriptional reporters were grown either in TBK-Brij or TBK-Ole. Cultures were harvested in the stationary phase, and β-gal activity was measured. For each reporter fusion, data were normalized to the β-gal activity of WT in TBK-Brij. Data represent the average (± SD) of three independent experiments. The average β-gal activity (in Miller units) of various WT reporter strains in TBK-Brij was: *fadE1000-lacZ* (58±5), *fadE400-lacZ* (88±21), and *fadE200-lacZ* (59±3). *B)* Similar to *fadE1000-lacZ*, *fadE200-lacZ* is also regulated by Cpx only in the stationary phase. Strains carrying *fadE200-lacZ* transcriptional reporter were grown either in TBK-Brij or TBK-Ole. Cultures were harvested in the exponential and stationary phases of growth. β-gal activity was measured, and data were normalized to the β-gal activity of WT grown in TBK-Brij at respective phases. Data represent the average (± SD) of three independent experiments. The average β-gal activity (in Miller units) of WT in TBK-Brij at exponential and stationary phases was 69±5 and 68±12, respectively. For panels A and B, the p-values were calculated using the unpaired two-tailed Student's t test (***, $P<0.001$; **, $P<0.01$; *, $P<0.05$; ns, $P>0.05$).

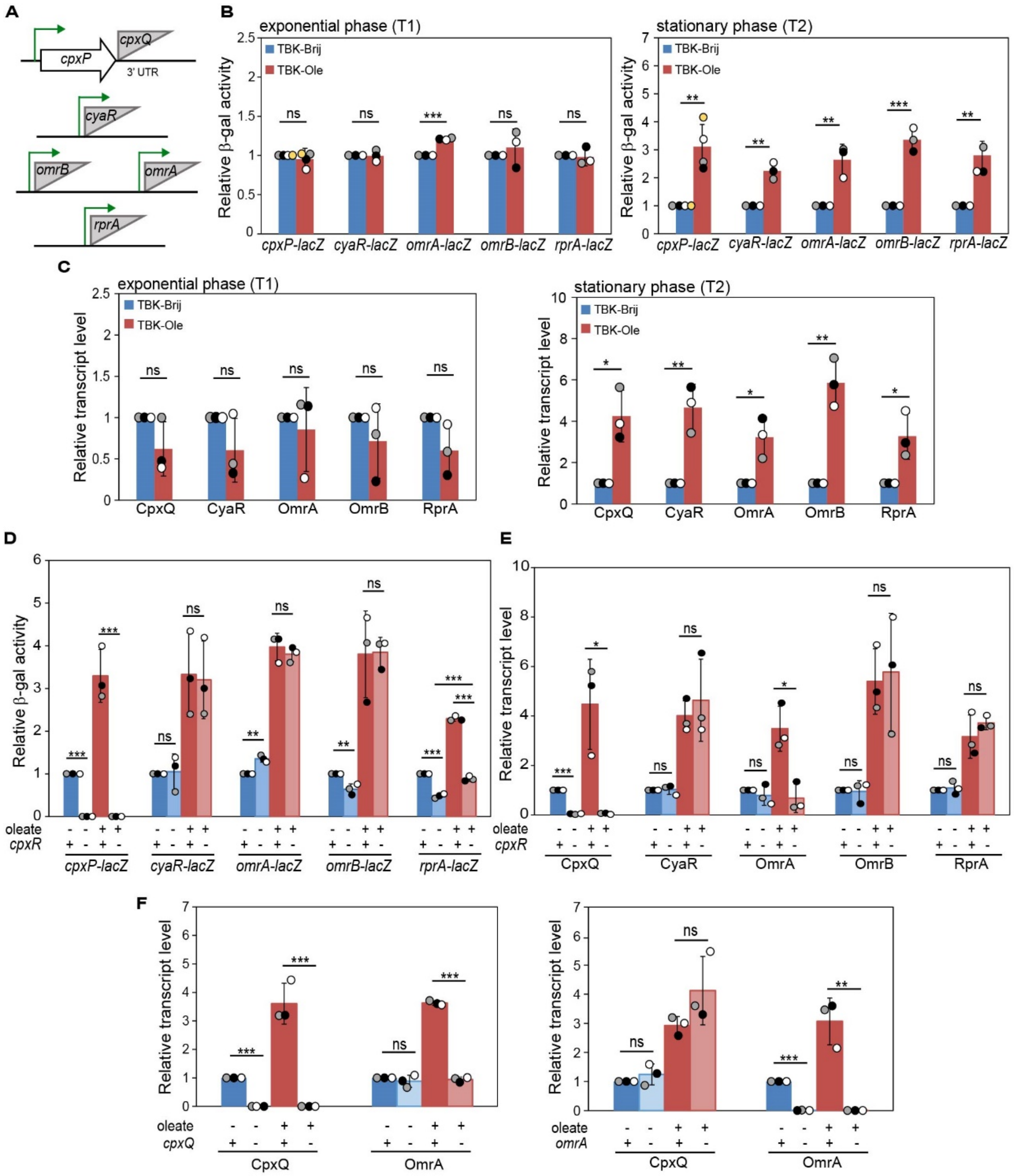

**Fig. S6.** *Of the five known Cpx-regulated sRNAs, in LCFA-grown cells, CpxR induces the expression of only CpxQ and increases the stability of OmrA via CpxQ. A)* Schematic of the genomic context of known Cpx-regulated sRNAs. Except for CpxQ, which is generated by RNase E cleavage of the 3' untranslated region (UTR) of *cpxP* mRNA, other sRNAs are expressed from their own promoter (1-4). *B)* All five sRNAs are transcriptionally induced in oleate-grown cells in the stationary phase. WT carrying chromosomal fusion of *lacZ* with the promoter of sRNAs was grown either in TBK-Brij or TBK-Ole. Cultures were harvested in exponential and stationary phases, and β-gal activity was measured. For each reporter fusion, data were normalized to the β-gal activity of WT in TBK-Brij at respective phases and represent the average (± SD) of at least three independent experiments. The average β-gal activity (in Miller units) of various reporter strains in TBK-Brij in the exponential phase was: *cpxP-lacZ* (810±109), *cyaR-lacZ* (308±7), *omrA-lacZ* (19±1), *omrB-lacZ* (79±12), and *rprA-lacZ* (463±167), and in the stationary phase was: *cpxP-lacZ* (311±147), *cyaR-lacZ* (183±33), *omrA-lacZ* (46±6), *omrB-lacZ* (65±11), and *rprA-lacZ* (190±17). *C)* The transcript levels of all five sRNAs increase in oleate-grown cells in the stationary phase. WT was grown either in TBK-Brij or TBK-Ole. Cultures were harvested for RNA isolation in exponential and stationary phases, and processed for qRT-PCR as mentioned in the legend to Fig 2A. Data were normalized to the transcript levels in WT in TBK-Brij at respective phases and represent the average (± SD) of three independent experiments. *D)* The transcriptional induction of CpxQ in oleate-grown cells is dependent on CpxR. Strains carrying chromosomal fusion of *lacZ* with the promoter of sRNAs were grown either in TBK-Brij or TBK-Ole. Cultures were harvested in the stationary phase, and β-gal activity was measured. For each reporter fusion, data were normalized to the β-gal activity of WT in TBK-Brij and represent the average (± SD) of three independent experiments. The average β-gal activity (in Miller units) of various WT reporter strains in TBK-Brij was: *cpxP*-*lacZ* (308±29), *cyaR*-*lacZ* (167±50), *omrA*-*lacZ* (50±4), *omrB*-*lacZ* (163±57), and *rprA*-*lacZ* (207±30). *E)* The transcript levels of CpxQ and OmrA increase in oleate-grown cells in a CpxR-dependent manner. Strains were grown either in TBK-Brij or TBK-Ole. Cultures were harvested in the stationary phase and processed for qRT-PCR as mentioned in the legend to Fig 2A. Data were normalized to the transcript levels in WT in TBK-Brij and represent the average (± SD) of three independent experiments. *F)* The increase in transcript levels of OmrA in oleate-grown cells is CpxQ-dependent. Strains were grown either in TBK-Brij or TBK-Ole till the stationary phase. Cultures were harvested and processed for qRT-PCR as mentioned in the legend to Fig 2A. Data were normalized to the transcript levels in WT in TBK-Brij and represent the average (± SD) of three independent experiments. For panels B-F, the p-values were calculated using the unpaired two-tailed Student's t test (***, P<0.001; **, P<0.01; *, P<0.05; ns, P>0.05).

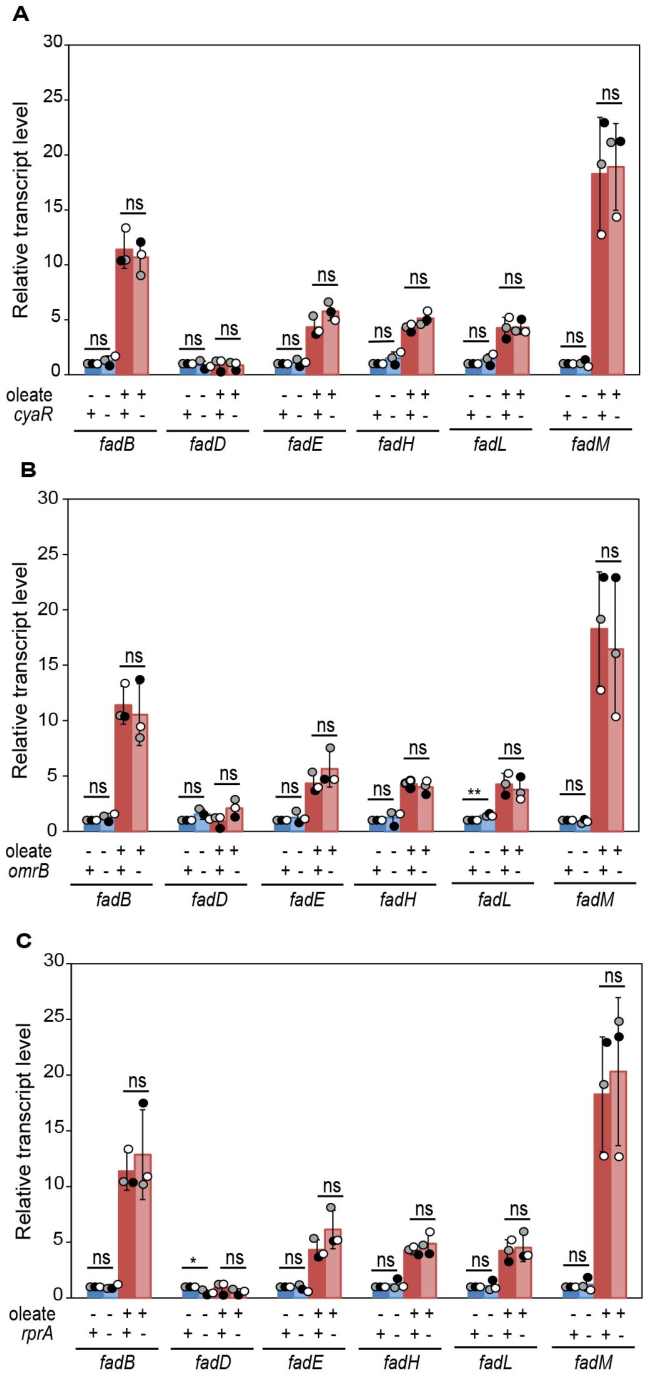

**Fig. S7.** *CyaR (A)*, *OmrB (B)*, *and RprA (C) do not regulate fad genes in LCFA-grown cells.* Strains were grown either in TBK-Brij or TBK-Ole. Cultures were harvested for RNA isolation in the stationary phase and processed for qRT-PCR, as mentioned in the legend to Fig 2A. Data were normalized to the transcript levels in WT in TBK-Brij and represent the average (± SD) of three independent experiments. In all three panels, the data shown for the fold change of transcript levels of various *fad* genes in the WT strain are the same. The p-values were calculated using the unpaired two-tailed Student's t test (***, P<0.001; **, P<0.01; *, P<0.05; ns, P>0.05).

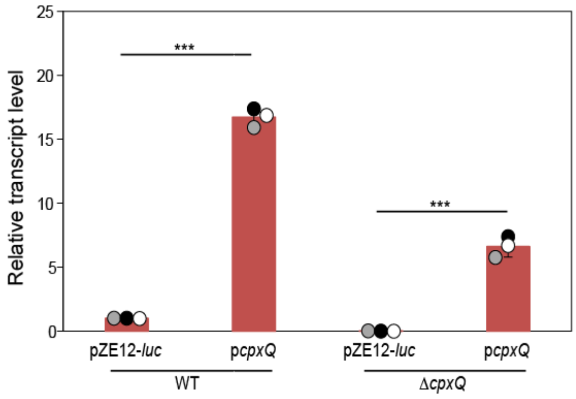

**Fig. S8.** *CpxQ is expressed from pZE12-luc*. Strains transformed with either pZE12-*luc* or p*cpxQ* were grown in TBK-Ole. Cultures were harvested for RNA isolation in the stationary phase and processed for qRT-PCR, as mentioned in the legend to Fig 2A. Data were normalized to the transcript levels of CpxQ in WT transformed with pZE12-*luc* and represent the average (± SD) of three independent experiments. The p-values were calculated using the unpaired two-tailed Student's t test (***, P<0.001).

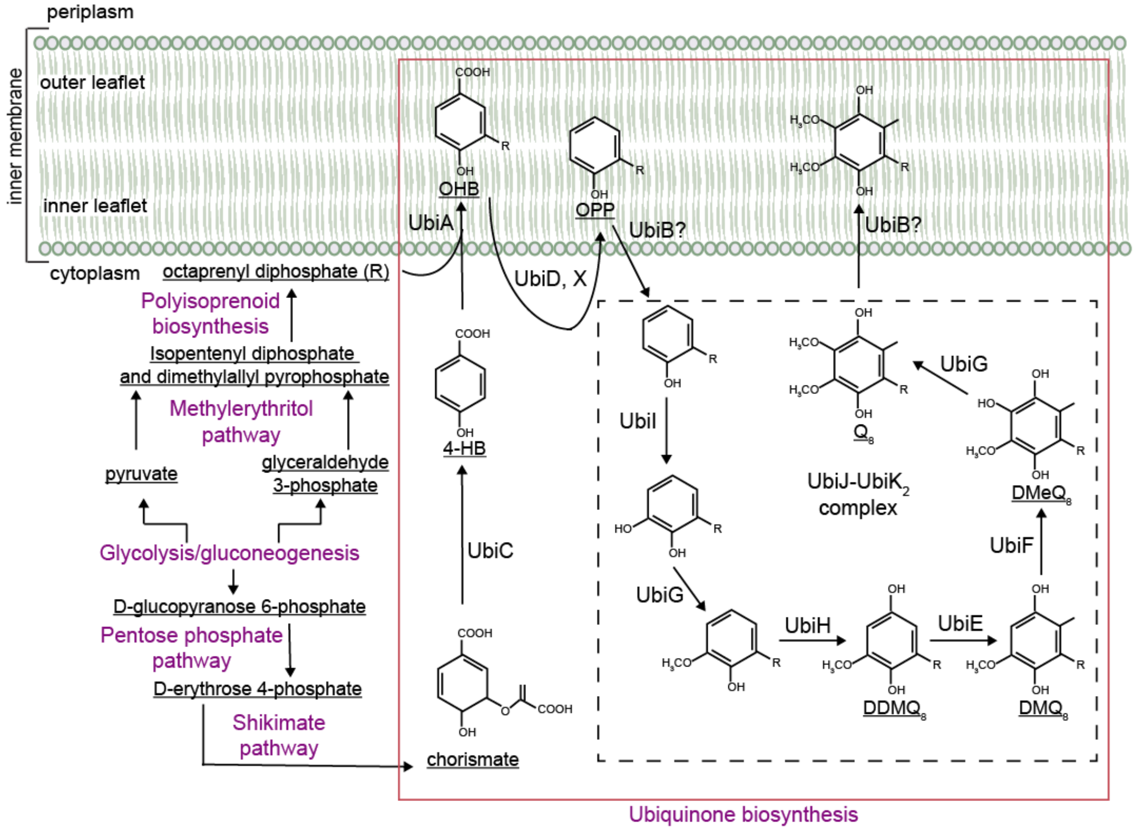

**Fig. S9.** *Ubiquinone biosynthesis in E. coli*. Glycolysis/gluconeogenesis provides the building blocks for the generation of ubiquinone precursors, chorismate and octaprenyl diphosphate (R), via multiple metabolic pathways. Chorismate is formed from the shikimate pathway, which uses D-erythrose 4-phosphate, a product of the pentose phosphate pathway, as a substrate. During growth on non-glycolytic substrates such as LCFAs, gluconeogenesis provides the precursor for D-erythrose 4-phosphate. Octaprenyl diphosphate is formed from the polyisoprenoid biosynthesis pathway which uses isopentenyl diphosphate (IPP) and dimethylallyl pyrophosphate (DMAPP) as substrates. IPP and DMAPP are the products of the methylerythritol phosphate (MEP) pathway which uses pyruvate and D-glyceraldehyde 3-phosphate, generated from gluconeogenesis (during growth on non-glycolytic substrates), as precursors. In *E. coli* at least twelve genes are involved in core ubiquinone biosynthesis (*ubiA* to *ubiK* and *ubiX*, shown within the red box) under aerobic conditions, most of them encode enzymes that modify the aromatic 4-hydroxybenzoate (4-HB) ring, which is formed by the removal of pyruvate from chorismate by UbiC (5-7). The black dotted rectangle delimits the Ubi-complex, composed of UbiE to UbiK proteins. Abbreviations: 4-HB, 4-hydroxybenzoate; OHB, octaprenyl-4-hydroxybenzoate; OPP, octaprenyl phenol; DDMQ8, C2-demethyl-C6-demethoxy-ubiquinone-8; DMQ8, C6-demethoxy-ubiquinone-8; DMeQ8, 6-demethyl-ubiquinone-8; Q8, ubiquinone-8.

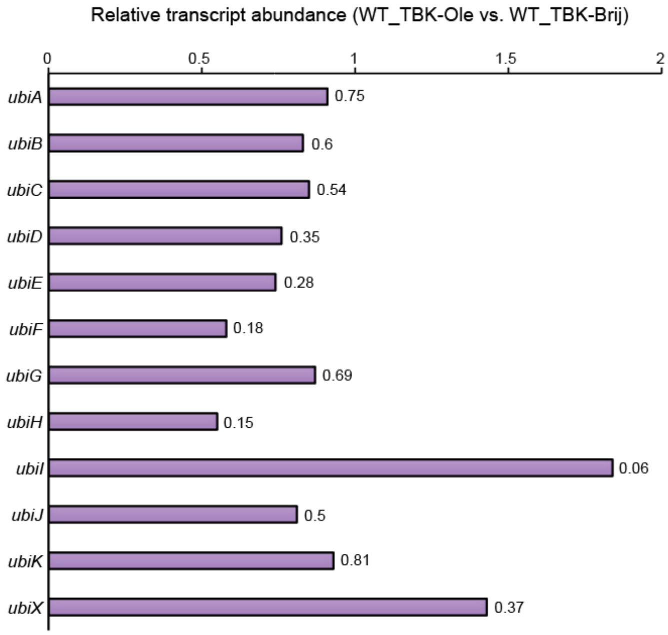

**Fig. S10.** *Relative abundance of ubi transcripts obtained from differential expression analysis*. Fold-change in *ubi* transcripts of the comparison group WT_TBK-Ole vs. WT_TBK-Brij obtained from differential expression analysis (Supplementary dataset S1) is plotted. Values indicated above each bar represent the p-value.

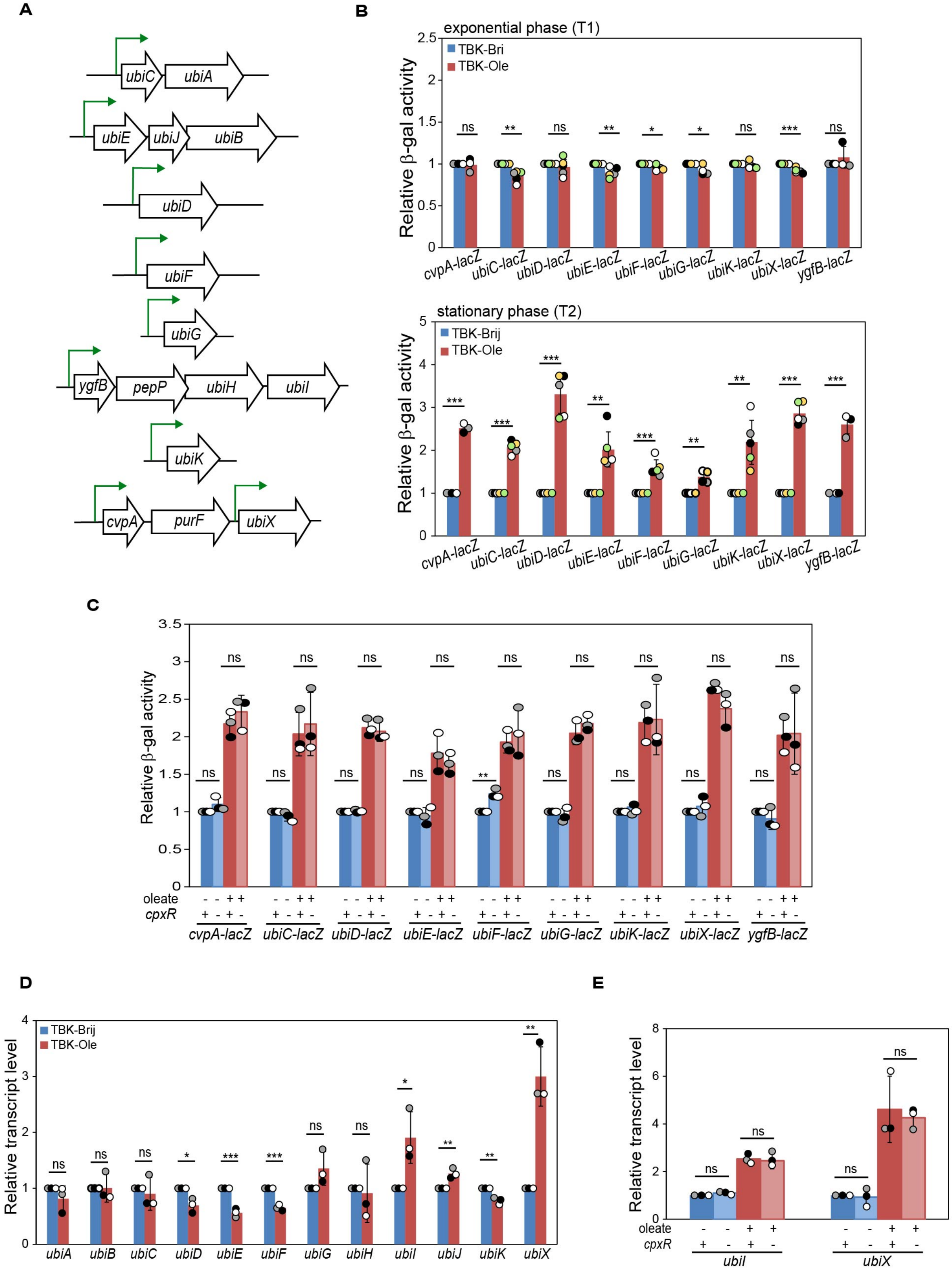

**Fig. S11.** *Several ubiquinone biosynthesis genes are upregulated in LCFA-grown cells; however, their upregulation is CpxR-independent. A)* Genetic organization of the *ubi* genes in *E. coli*. Some *ubi* genes are present alone on the chromosome while others are part of an operon (6). To cover the possibility of internal promoters that might be active in the LCFA condition, besides making chromosomal *lacZ* reporter fusions with the *cis*-acting element of the first gene of the operon, we also made reporter fusions with sequences immediately upstream of each *ubi* gene located within the operon. Whereas all reporter strains carrying the *cis*-acting element of the first gene of the operon showed higher reporter expression than the strain carrying promoterless reporter; except *ubiX*, reporter strains harboring sequences upstream of the *ubi* genes within the operon exhibited near background reporter expression (Table S2). Green arrows denote transcription from the active promoters. *B) ubi* genes are upregulated in the stationary phase in oleate-grown cells. WT carrying chromosomal *lacZ* transcriptional reporters was grown either in TBK-Brij or TBK-Ole. Cultures were harvested in exponential (upper panel) and stationary (lower panel) phases, and β-gal activity was measured. For each reporter fusion, data were normalized to the β-gal activity of WT in TBK-Brij and represent the average (± SD) of at least three independent experiments. The average β-gal activity (in Miller units) of various reporter strains is provided in Table S2. *C)* The upregulation of *ubi* genes in oleate-grown cells is CpxR-independent. Strains carrying chromosomal *lacZ* transcriptional reporters were grown either in TBK-Brij or TBK-Ole. Cultures were harvested in the stationary phase, and β-gal activity was measured. For each reporter fusion, data were normalized to the β-gal activity of WT in TBK-Brij and represent the average (± SD) of three independent experiments. The average β-gal activity (in Miller units) of various reporter strains in TBK-Brij was: *cvpA-lacZ* (58±5), *ubiC-lacZ* (56±6), *ubiD-lacZ* (84±6), *ubiE-lacZ* (162±13), *ubiF-lacZ* (57±5), *ubiG-lacZ* (91±4), *ubiK-lacZ* (172±33), *ubiX-lacZ* (26±2), and *ygfB-lacZ* (66±11). *D & E)* Amongst the ubiquinone biosynthesis genes, only the transcript levels of *ubiI* and *ubiX* increase in LCFA-grown cells *(D)*; however, their increase is CpxR-independent *(E).* Strains were grown either in TBK-Brij or TBK-Ole till the stationary phase. Cultures were harvested for RNA isolation in the stationary phase and processed for qRT-PCR, as mentioned in the legend to Fig 2A. Data were normalized to the transcript levels in WT in TBK-Brij and represent the average (± SD) of three independent experiments. For panels B-E, the p-values were calculated using the unpaired two-tailed Student's t test (***, P<0.001; **, P<0.01; *, P<0.05; ns, P>0.05).

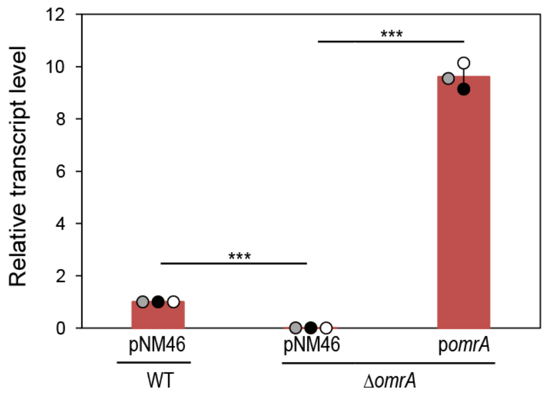

**Fig. S12.** *OmrA is expressed from pNM46*. Strains transformed with either pNM46 or p*omrA* were grown in TBK-Ole supplemented with 0.1 mM IPTG. Cultures were harvested for RNA isolation in the stationary phase and processed for qRT-PCR, as mentioned in the legend to Fig 2A. Data were normalized to the transcript levels of OmrA in WT transformed with pNM46 and represent the average (± SD) of three independent experiments. The p-values were calculated using the unpaired two-tailed Student's t test (***, P<0.001).

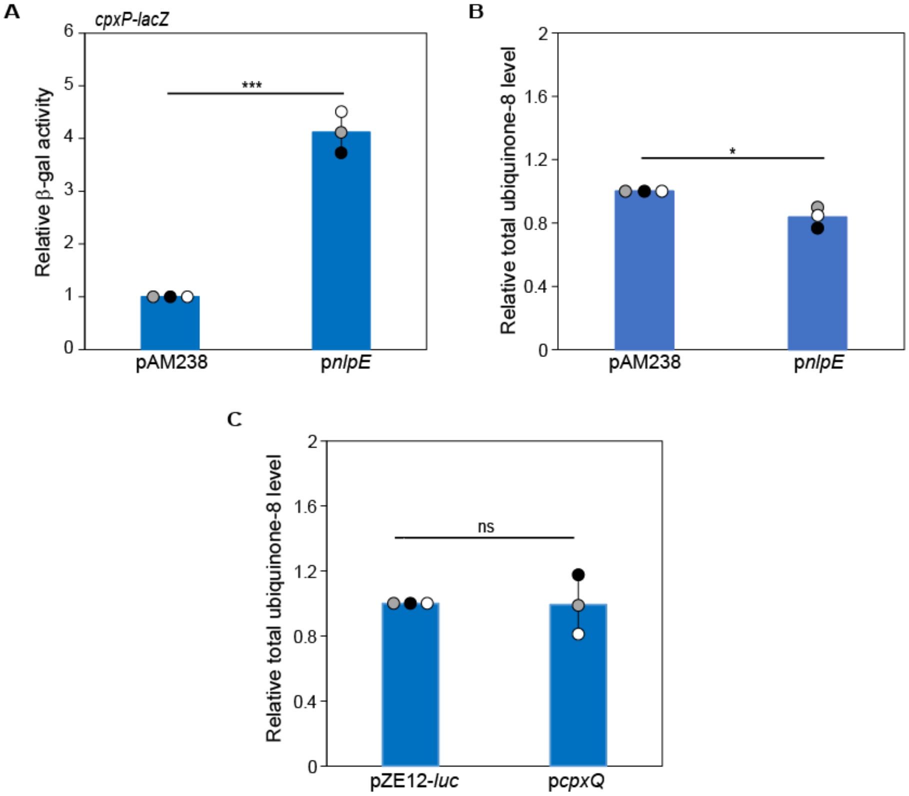

**Fig. S13.** *Activation of the Cpx pathway or CpxQ overexpression is not sufficient to increase ubiquinone levels. A)* NlpE overexpression induces *cpxP*. The *cpxP-lacZ* reporter strain was transformed with either pAM238 or pAM238 carrying *nlpE* (p*nlpE*) and was grown in TBK medium supplemented with 0.1 mM IPTG. Cultures were harvested in the exponential phase, and β-gal activity was measured. Data were normalized to the β-gal activity of WT carrying pAM238 and represent the average (± SD) of three independent experiments. The average β-gal activity (in Miller units) of WT carrying pAM238 was 646±55. *B)* NlpE overexpression does not alter ubiquinone levels. WT transformed with either pAM238 or p*nlpE* was grown in TBK medium supplemented with 0.1 mM IPTG. The total ubiquinone-8 content in the lipid extracts was measured in the exponential phase. The total ubiquinone-8 level in each sample was normalized to the total ubiquinone-8 level of WT transformed with pAM238. Data represent the average (± SD) of three independent experiments. *C*) Ubiquinone levels remain unchanged upon CpxQ overexpression. WT transformed with either pZE12-*luc* or p*cpxQ* was grown in TBK-Brij. The total ubiquinone-8 content in the lipid extracts was measured in the stationary phase. The total ubiquinone-8 level in each sample was normalized to the total ubiquinone-8 level of WT transformed with pZE12-*luc*. Data represent the average (± SD) of three independent experiments. The p-values were calculated using the unpaired two-tailed Student's t test (***, $P<0.001$; **, $P<0.01$; *, $P<0.05$; ns, $P>0.05$).

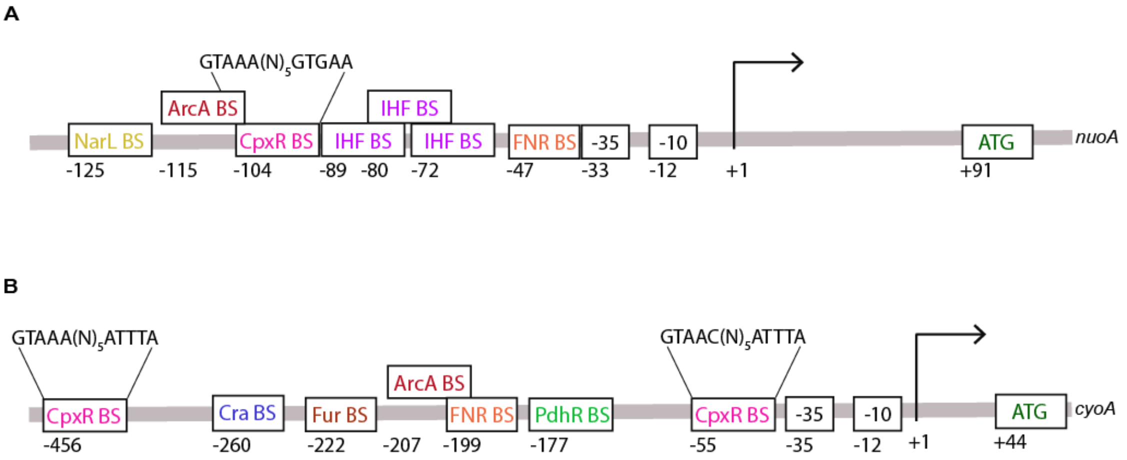

**Fig. S14.** *Identification of a putative CpxR binding site upstream of nuoA and cyoA using Virtual Footprint.* Schematic representation of the *nuoA* (A) and *cyoA* (B) promoter region of *E. coli* K-12 MG1655 indicating the location of the binding sites for CpxR (obtained from Virtual Footprint (8)) and other transcription factors (obtained from EcoCyc (9)). Numbers indicate distance from the transcription start site (+1), where '-' and '+' denote upstream and downstream, respectively. Abbreviation: BS, binding site.

**Table S1.** Normalized counts for genes constituting LCFA-related pathways used to generate Fig. 1A

| Pathway | Gene name | WT_ TBK-Brij | Δ*cpxR*_ TBK-Brij | WT_ TBK-Ole | Δ*cpxR*_ TBK-Ole |
|---|---|---|---|---|---|
| LCFA transport and β-oxidation | *fadA* | 1027.136905 | 908.3797895 | 4537.286497 | 9764.003978 |
| | *fadB* | 2049.845078 | 2070.888748 | 11987.32024 | 24640.91236 |
| | *fadD* | 2662.951326 | 2895.669716 | 2970.680977 | 4047.620829 |
| | *fadE* | 5486.069375 | 4598.562647 | 17943.98572 | 31368.66725 |
| | *fadH* | 966.9827974 | 921.2801929 | 2432.631322 | 4367.152538 |
| | *fadI* | 653.9326933 | 649.8167643 | 1890.741275 | 3250.800547 |
| | *fadJ* | 797.3428056 | 768.2084468 | 2181.885034 | 4245.12103 |
| | *fadL* | 1555.68853 | 1482.236949 | 3034.221058 | 6187.337108 |
| | *fadM* | 135.9200337 | 115.1307153 | 529.0740985 | 1006.387286 |
| TCA cycle | *acnA* | 15958.1858 | 12666.01838 | 6869.680045 | 2904.421181 |
| | *acnB* | 2324.043133 | 1593.511988 | 4978.156596 | 8757.220143 |
| | *fumA* | 562.177921 | 637.85659 | 914.1422979 | 1417.105808 |
| | *fumB* | 42.07870867 | 55.06503723 | 82.20282952 | 68.7010528 |
| | *fumC* | 943.1590075 | 1240.678569 | 569.0416855 | 767.7291088 |
| | *gltA* | 5668.80767 | 5516.997566 | 6242.129963 | 8323.679462 |
| | *icd* | 986.3962884 | 1147.3062 | 2482.446103 | 6118.575291 |
| | *lpd* | 1041.345675 | 953.3741405 | 2282.384978 | 3947.7493 |
| | *maeA* | 582.1675793 | 612.6597957 | 1063.472397 | 1905.528647 |
| | *mdh* | 3490.684996 | 3112.140665 | 3027.998498 | 3832.263301 |
| | *mqo* | 257.0269231 | 196.2736306 | 200.0819275 | 129.3862483 |
| | *sdhA* | 2010.328179 | 2996.65538 | 3224.805941 | 4244.223267 |
| | *sdhB* | 458.0013738 | 690.2007792 | 851.4541724 | 1325.712868 |
| | *sdhC* | 848.4151896 | 1277.225623 | 1025.337899 | 1200.623636 |
| | *sdhD* | 441.2036081 | 675.799047 | 657.9385836 | 834.7651712 |
| | *sucA* | 3787.121209 | 4953.111306 | 6215.327963 | 8156.699817 |
| | *sucB* | 2025.666145 | 2762.095153 | 3431.165927 | 4874.293943 |
| | *sucC* | 1651.837864 | 2251.202923 | 3053.65565 | 5011.779837 |
| | *sucD* | 1218.816326 | 1564.469057 | 2256.975539 | 3606.918027 |
| Glyoxylate shunt | *aceA* | 16384.31317 | 15656.40836 | 26116.47425 | 49690.24408 |
| | *aceB* | 16429.5464 | 14833.04528 | 19743.33196 | 32132.13384 |
| | *acnA* | 15958.1858 | 12666.01838 | 6869.680045 | 2904.421181 |
| | *acnB* | 2324.043133 | 1593.511988 | 4978.156596 | 8757.220143 |
| | *gltA* | 5668.80767 | 5516.997566 | 6242.129963 | 8323.679462 |
| | *maeA* | 582.1675793 | 612.6597957 | 1063.472397 | 1905.528647 |
| | *mdh* | 3490.684996 | 3112.140665 | 3027.998498 | 3832.263301 |
| Gluconeogenesis | *eno* | 7753.476794 | 6037.893199 | 4536.346387 | 3685.426227 |
| | *fbaA* | 2576.905958 | 2061.35265 | 2059.49722 | 2503.432245 |
| | *fbaB* | 5511.946003 | 5367.854851 | 3427.174677 | 2850.862634 |
| | *fbp* | 304.8302945 | 361.8675707 | 359.5119472 | 555.123777 |
| | *gapA* | 6008.858977 | 6128.008142 | 5285.94443 | 5384.606535 |
| | *glpX* | 444.8132972 | 379.5412627 | 225.9774587 | 212.3408602 |
| | *gpmM* | 616.3791511 | 545.973578 | 489.3322662 | 382.1625509 |
| | *kduI* | 54.22912756 | 340.2737693 | 54.54703185 | 53.94980158 |
| | *maeA* | 582.1675793 | 612.6597957 | 1063.472397 | 1905.528647 |
| | *maeB* | 666.6619042 | 656.1148165 | 1100.966839 | 2067.508505 |

| | | | | | |
|---|---|---|---|---|---|
| | *mdh* | 3490.684996 | 3112.140665 | 3027.998498 | 3832.263301 |
| | *pck* | 802.4642929 | 1020.950616 | 1482.43327 | 2569.345129 |
| | *pgi* | 2728.172827 | 2831.742704 | 2002.550112 | 1554.265288 |
| | *pgk* | 3782.15838 | 3144.317706 | 2840.114652 | 2918.710801 |
| | *ppsA* | 869.4173889 | 1021.40557 | 1771.329474 | 4532.152678 |
| | *tpiA* | 1647.716479 | 1579.448172 | 1343.6678 | 1604.005889 |
| | *ybhA* | 49.41624473 | 64.36602995 | 82.42972394 | 102.1497169 |
| | *ybiV* | 112.6236496 | 83.14301145 | 98.02959937 | 103.5562089 |
| | *yggF* | 62.72625531 | 46.11712194 | 68.3637698 | 47.0533688 |
| | *yidA* | 66.80773932 | 57.15040585 | 102.8629916 | 139.5896869 |
| | *yigL* | 127.3114673 | 128.7918135 | 153.2061097 | 147.1901042 |
| NADH dehydrogenase I | *nuoA* | 330.1385488 | 489.8952367 | 412.9116676 | 691.5732125 |
| | *nuoB* | 508.337146 | 676.7804391 | 616.4533026 | 1164.594589 |
| | *nuoC* | 906.8502332 | 1122.833416 | 1183.344915 | 2035.075367 |
| | *nuoE* | 262.4317458 | 337.2951264 | 353.0629093 | 753.0403976 |
| | *nuoF* | 507.2468678 | 624.5007223 | 707.5107528 | 1419.076972 |
| | *nuoG* | 1672.655149 | 2171.013282 | 1981.299339 | 3682.296252 |
| | *nuoH* | 304.4640698 | 498.9518301 | 457.6692046 | 853.4757164 |
| | *nuoI* | 192.5423475 | 271.4939663 | 246.0582857 | 513.3939364 |
| | *nuoJ* | 251.9700858 | 347.1794545 | 351.1306918 | 656.9631673 |
| | *nuoK* | 112.6158414 | 147.2857211 | 156.1068988 | 304.2039336 |
| | *nuoL* | 580.4221901 | 862.0519932 | 895.774779 | 1718.466039 |
| | *nuoM* | 158.4622427 | 266.8001919 | 434.3915579 | 879.4308223 |
| | *nuoN* | 340.0361976 | 493.0225784 | 563.9595748 | 999.9364588 |
| Cytochrome *bo* oxidase | *cyoA* | 326.2684308 | 410.1145633 | 857.2301791 | 1703.255363 |
| | *cyoB* | 398.3643083 | 596.091562 | 1085.310702 | 2267.422394 |
| | *cyoC* | 106.1745312 | 137.36492 | 279.4497775 | 520.9346955 |
| | *cyoD* | 105.8930509 | 111.9707783 | 229.8930201 | 419.6146385 |
| ATP biosynthesis | *atpA* | 1216.853306 | 1029.326858 | 3071.288718 | 5810.576569 |
| | *atpB* | 408.5879981 | 421.8103794 | 839.2329963 | 1395.890467 |
| | *atpC* | 251.7057942 | 288.5452617 | 655.0231251 | 1263.760298 |
| | *atpD* | 758.9257199 | 736.2117826 | 2008.139149 | 3998.942199 |
| | *atpE* | 278.9961094 | 266.7018542 | 741.0534667 | 1378.655533 |
| | *atpF* | 484.3902779 | 390.2202203 | 1225.226527 | 2294.417273 |
| | *atpG* | 465.5478679 | 398.302958 | 1182.433232 | 2379.459371 |
| | *atpH* | 232.3482557 | 222.3083249 | 551.7324375 | 1148.019975 |

**Table S2.** Miller units (MU) of various transcriptional reporter strains in exponential (exp) and stationary (st) phases

| Transcriptional reporter | MU in exp phase (average ± SD) | | MU in st phase (average ± SD) | |
|---|---|---|---|---|
| | TBK-Brij | TBK-Ole | TBK-Brij | TBK-Ole |
| promoterless *lacZ* | 11.0±3.9 | 9.5±4.8 | 18.6±4.5 | 21.5±4.8 |
| *cvpA-lacZ* | 60.3±6.2 | 59.8±9.2 | 52.0±1.5 | 131.0±2.8 |
| *ubiA-lacZ* | 7.2±4.6 | 5.2±2.4 | 5.8±1.3 | 9.3±3.0 |
| *ubiB-lacZ* | 5.5±1.4 | 7.4±0.6 | 18.8±5.1 | 21.7±8.0 |
| *ubiC-lacZ* | 68.6±20.9 | 57.6±12.4 | 46.3±8.1 | 95.8±18.6 |
| *ubiD-lacZ* | 94.5±10.2 | 90.4±7.0 | 38.6±6.4 | 125.8±14.6 |
| *ubiE-lacZ* | 181.0±24.9 | 161.7±17.2 | 131.7±25.2 | 260.4±42.9 |
| *ubiF-lacZ* | 64.5±18.0 | 61.4±16.4 | 69.1±13.8 | 108.5±13.8 |
| *ubiG-lacZ* | 240.8±90.4 | 219.4±74.2 | 127.7±20.5 | 178.9±46.3 |
| *ubiH-lacZ* | 4.2±1.3 | 3.7±0.8 | 6.2±2.2 | 6.4±1.0 |
| *ubiI-lacZ* | 16.1±7.4 | 14.8±7.3 | 13.6±1.1 | 17.3±1.5 |
| *ubiJ-lacZ* | 11.5±2.0 | 9.7±1.8 | 29.0±2.9 | 34.3±1.9 |
| *ubiK-lacZ* | 220.6±15.5 | 217.0±17.4 | 125.4±46.4 | 257.6±82.8 |
| *ubiX-lacZ* | 55.5±19.8 | 50.8±17.4 | 17.9±6.2 | 50.6±15.6 |
| *ygfB-lacZ* | 65.1±9.2 | 70.8±18.1 | 50.9±5.1 | 131.6±4.5 |

* The transcriptional reporter strains that showed higher reporter expression than the promoterless reporter strain are highlighted.

**Table S3.** Strains and Plasmids used in this study

| Strains/Plasmids | Relevant genotype | Source (Reference) |
|---|---|---|
| **Strains** | | |
| BW25113 | $F^-$ Δ(*araD-araB*)*567* Δ*lacZ4787*(::rrnB-3) $\lambda^-$ *rph-1* Δ(*rhaD-rhaB*)*568 hsdR514* | *E. coli* Genetic Stock Centre (10) |
| BW25142 | *lacI$^q$ rrnB3* Δ*lacZ4787 hsdR514* Δ(*araBAD*)*567* Δ(*rhaBAD*)*568* Δ*phoBR580 rph-1 galU95* Δ*endA9 uidA*(Δ*MluI*)::*pir-116 recA1* | Rao lab (11) |
| DH5α | $F^-$ Δ(*argF-lac*)*169* φ80d*lacZ58*(M15) *glnX44*(AS) $\lambda^-$ *rfbC1 gyrA96*(Nal$^r$) *recA1 endA1 spoT1 thiE1 hsdR17* | New England Biolabs |
| *cpxR*::*kan* | BW25113 *cpxR*::*kan*, Kan$^r$ | Keio collection (10) |
| *hfq*::*kan* | BW25113 *hfq*::*kan*, Kan$^r$ | Keio collection (10) |
| *cpxR*::*cam* | BW25113 *cpxR*::*cam*, Cam$^r$ | ASKA library (12) |
| *fadR*::*cam* | BW25113 *fadR*::*cam*, Cam$^r$ | ASKA library (12) |
| RC22026 | BW25113 *cpxQ*::*kan*, Kan$^r$ | This work |
| RC22081 | BW25113 *cpxQ::cam*, Cam$^r$ | This work |
| RC19096 | BW25113 *cyaR*::*kan*, Kan$^r$ | This work |
| RC19094 | BW25113 *omrA*::*kan*, Kan$^r$ | This work |
| RC19095 | BW25113 *omrB*::*kan*, Kan$^r$ | This work |
| RC22012 | BW25113 *rprA*::*kan*, Kan$^r$ | This work |
| RC15087 (promoterless-*lacZ*) | BW25113 *att*λ::[Kan promoterless *pAH125 oriR6K*], Kan$^r$ | This work |
| RC18098 (*cpxP-lacZ*) | BW25113 *att*λ::[Kan $P_{cpxP(-410/+223)}$-*lacZ oriR6K*], Kan$^r$ | This work |
| Freshly made | P1 (BW25113 *cpxR*::*cam*) x RC18098, Kan$^r$ Cam$^r$ | This work |
| RC15082 (*fadE*$_{1000}$-*lacZ*) | BW25113 *att*λ::[Kan $P_{fadE1000}$-*lacZ oriR6K*], Kan$^r$ | (13) |
| Freshly made | P1 (BW25113 *cpxR*::*cam*) x RC15082, Kan$^r$ Cam$^r$ | This work |
| RC21022 (*fadE*$_{400}$-*lacZ*) | BW25113 *att*λ::[Kan $P_{fadE400}$-*lacZ oriR6K*], Kan$^r$ | This work |
| Freshly made | P1 (BW25113 *cpxR*::*cam*) x RC21022, Kan$^r$ Cam$^r$ | This work |
| RC21023 (*fadE*$_{200}$-*lacZ*) | BW25113 *att*λ::[Kan $P_{fadE200}$-*lacZ oriR6K*], Kan$^r$ | This work |
| Freshly made | P1 (BW25113 *cpxR*::*cam*) x RC21023, Kan$^r$ Cam$^r$ | This work |
| Freshly made | P1 (BW25113 *cpxQ*::*cam*) x RC21023, Kan$^r$ Cam$^r$ | This work |
| RC22050 | P1 (BW25113 *fadR*::*cam*) x RC21023, Kan$^r$ Cam$^r$ | This work |
| RC22051 | Δ*fadR* RC21023 (Cam cassette flipped from RC22050), Kan$^r$ | This work |
| Freshly made | P1 (BW25113 *cpxR*::*cam*) x RC22051, Kan$^r$ Cam$^r$ | This work |
| Freshly made | P1 (BW25113 *cpxQ*::*cam*) x RC22051, Kan$^r$ Cam$^r$ | This work |
| RC15171 (*fadB*$_{200}$-*lacZ*) | BW25113 *att*λ::[Kan $P_{fadB200}$-*lacZ oriR6K*], Kan$^r$ | This work |
| Freshly made | P1 (BW25113 *cpxR*::*cam*) x RC15171, Kan$^r$ Cam$^r$ | This work |
| RC28010 (*fadD*$_{200}$-*lacZ*) | BW25113 *att*λ::[Kan $P_{fadD200}$-*lacZ oriR6K*], Kan$^r$ | This work |
| Freshly made | P1 (BW25113 *cpxR*::*cam*) x RC28010, Kan$^r$ Cam$^r$ | This work |
| Freshly made | P1 (BW25113 *cpxQ*::*cam*) x RC28010, Kan$^r$ Cam$^r$ | This work |
| RC15188 (*fadH*$_{200}$-*lacZ*) | BW25113 *att*λ::[Kan $P_{fadH200}$-*lacZ oriR6K*], Kan$^r$ | This work |
| Freshly made | P1 (BW25113 *cpxR*::*cam*) x RC15188, Kan$^r$ Cam$^r$ | This work |

| | | |
|---|---|---|
| RC22093 (*fadL*$_{200}$-*lacZ*) | BW25113 *att*λ::[Kan $P_{fadL200}$-*lacZ oriR6K*], $Kan^r$ | This work |
| Freshly made | P1 (BW25113 *cpxR*::*cam*) x RC22093, $Kan^r$ $Cam^r$ | This work |
| RC15169 (*fadM*$_{200}$-*lacZ*) | BW25113 attλ::[Kan $P_{fadM200}$-*lacZ oriR6K*], $Kan^r$ | This work |
| Freshly made | P1 (BW25113 *cpxR*::*cam*) x RC15169, $Kan^r$ $Cam^r$ | This work |
| RC22094 (*fadR*$_{200}$-*lacZ*) | BW25113 *att*λ::[Kan $P_{fadR200}$-*lacZ oriR6K*], $Kan^r$ | This work |
| Freshly made | P1 (BW25113 *cpxR*::*cam*) x RC22094, $Kan^r$ $Cam^r$ | This work |
| RC19115 (*cyaR-lacZ*) | BW25113 *att*λ::[Kan $P_{cyaR200}$-*lacZ oriR6K*], $Kan^r$ | This work |
| Freshly made | P1 (BW25113 *cpxR*::*cam*) x RC19115, $Kan^r$ $Cam^r$ | This work |
| RC22045 (*omrA-lacZ*) | BW25113 *att*λ::[Kan $P_{omrA200}$-*lacZ oriR6K*], $Kan^r$ | This work |
| Freshly made | P1 (BW25113 *cpxR*::*cam*) x RC22045, $Kan^r$ $Cam^r$ | This work |
| RC15193 (*omrB-lacZ*) | BW25113 *att*λ::[Kan $P_{omrB200}$-*lacZ oriR6K*], $Kan^r$ | This work |
| Freshly made | P1 (BW25113 *cpxR*::*cam*) x RC15193, $Kan^r$ $Cam^r$ | This work |
| RC19117 (*rprA-lacZ*) | BW25113 *att*λ::[Kan $P_{rprA200}$-*lacZ oriR6K*], $Kan^r$ | This work |
| Freshly made | P1 (BW25113 *cpxR*::*cam*) x RC19117, $Kan^r$ $Cam^r$ | This work |
| RC15190 (*cvpA-lacZ*) | BW25113 *att*λ::[Kan $P_{cvpA350}$-*lacZ oriR6K*], $Kan^r$ | This work |
| Freshly made | P1 (BW25113 *cpxR*::*cam*) x RC15190, $Kan^r$ $Cam^r$ | This work |
| RC19086 (*ubiA-lacZ*) | BW25113 *att*λ::[Kan $P_{ubiA350}$-*lacZ oriR6K*], $Kan^r$ | This work |
| RC19041 (*ubiB-lacZ*) | BW25113 *att*λ::[Kan $P_{ubiB350}$-*lacZ oriR6K*], $Kan^r$ | This work |
| RC19048 (*ubiC-lacZ*) | BW25113 *att*λ::[Kan $P_{ubiC1000}$-*lacZ oriR6K*], $Kan^r$ | This work |
| Freshly made | P1 (BW25113 *cpxR*::*cam*) x RC19048, $Kan^r$ $Cam^r$ | This work |
| RC19055 (*ubiD-lacZ*) | BW25113 *att*λ::[Kan $P_{ubiD600}$-*lacZ oriR6K*], $Kan^r$ | This work |
| Freshly made | P1 (BW25113 *cpxR*::*cam*) x RC19055, $Kan^r$ $Cam^r$ | This work |
| RC19042 (*ubiE-lacZ*) | BW25113 *att*λ::[Kan $P_{ubiE350}$-*lacZ oriR6K*], $Kan^r$ | This work |
| Freshly made | P1 (BW25113 *cpxR*::*cam*) x RC19042, $Kan^r$ $Cam^r$ | This work |
| RC19043 (*ubiF-lacZ*) | BW25113 *att*λ::[Kan $P_{ubiF400}$-*lacZ oriR6K*], $Kan^r$ | This work |
| Freshly made | P1 (BW25113 *cpxR*::*cam*) x RC19043, $Kan^r$ $Cam^r$ | This work |
| RC19056 (*ubiG-lacZ*) | BW25113 *att*λ::[Kan $P_{ubiG350}$-*lacZ oriR6K*], $Kan^r$ | This work |
| Freshly made | P1 (BW25113 *cpxR*::*cam*) x RC19056, $Kan^r$ $Cam^r$ | This work |
| RC19006 (*ubiH-lacZ*) | BW25113 *att*λ::[Kan $P_{ubiH1000}$-*lacZ oriR6K*], $Kan^r$ | This work |
| RC19044 (*ubiI-lacZ*) | BW25113 *att*λ::[Kan $P_{ubiI350}$-*lacZ oriR6K*], $Kan^r$ | This work |
| RC19045 (*ubiJ-lacZ*) | BW25113 *att*λ::[Kan $P_{ubiJ350}$-*lacZ oriR6K*], $Kan^r$ | This work |
| RC19046 (*ubiK-lacZ*) | BW25113 *att*λ::[Kan $P_{ubiK350}$-*lacZ oriR6K*], $Kan^r$ | This work |
| Freshly made | P1 (BW25113 *cpxR*::*cam*) x RC19046, $Kan^r$ $Cam^r$ | This work |
| RC19051 (*ubiX-lacZ*) | BW25113 *att*λ::[Kan $P_{ubiX350}$-*lacZ oriR6K*], $Kan^r$ | This work |
| Freshly made | P1 (BW25113 *cpxR*::*cam*) x RC19051, $Kan^r$ $Cam^r$ | This work |

| | | |
|---|---|---|
| RC15191 (*ygfB-lacZ*) | BW25113 *att*λ::[Kan $P_{ygfB500}$-*lacZ oriR6K*], $Kan^r$ | This work |
| Freshly made | P1 (BW25113 *cpxR*::*cam*) x RC15191, $Kan^r$ $Cam^r$ | This work |
| RC22054 | BW25113 *fadR*-SPA, $Kan^r$ | This work |
| Freshly made | P1 (BW25113 *cpxQ*::*cam*) x RC22054, $Kan^r$ $Cam^r$ | This work |
| RC25029 | BW25113 *fadD*-SPA, $Kan^r$ | This work |
| Freshly made | P1 (BW25113 *cpxQ*::*cam*) x RC25029, $Kan^r$ $Cam^r$ | This work |
| RC25028 | BW25113 *fadL*-SPA, $Kan^r$ | This work |
| Freshly made | P1 (BW25113 *cpxQ*::*cam*) x RC25028, $Kan^r$ $Cam^r$ | This work |
| Freshly made | P1 (BW25113 *cpxR*::*cam*) x BW25113 *hfq*::*kan*, $Kan^r$ $Cam^r$ | This work |
| **Plasmids** | | |
| pKD13 | *oriR6K*, FRT-flanked $Kan^r$, pANTSγ PS1 PS4, $Kan^r$ | (14) |
| pKD3 | *oriR6K*, FRT-flanked $Cam^r$, pANTSγ PS1 PS4, $Amp^r$ $Cam^r$ | (14) |
| pSim5 | pSC101 *ori*, $P_L$-*gam-bet-exo* genes under the control of CI857 repressor (ts), $Cam^r$ | Court lab (15) |
| pSH06 | pSC101 *ori*, $P_L$-*gam-bet-exo* genes under the control of CI857 repressor (ts), $Amp^r$ | This work |
| pCP20 | pSC101 *ori*, *cI857* λ-$P_R$ *flp* ts, $Amp^r$ $Cam^r$ | (14) |
| pINT-ts | *oriR6K*, *int*, $Amp^r$ | Rao lab (11) |
| pAH125 | *oriR6K*, MCS-*lacZ t0 att*λ, $Kan^r$ | Rao lab (11) |
| pDR10 | *oriR6K*, MCS $P_{cpxP(-410/+223)}$-*lacZ t0 att*λ, $Kan^r$ | This work |
| pMS02 | *oriR6K*, MCS $P_{fadE1000}$-*lacZ t0 att*λ, $Kan^r$ | (13) |
| pSH02 | *oriR6K*, MCS $P_{fadE400}$-*lacZ t0 att*λ, $Kan^r$ | This work |
| pSH01 | *oriR6K*, MCS $P_{fadE200}$-*lacZ t0 att*λ, $Kan^r$ | This work |
| pMS13 | *oriR6K*, MCS $P_{fadB200}$-*lacZ t0 att*λ, $Kan^r$ | This work |
| pVS01 | *oriR6K*, MCS $P_{fadD200}$-*lacZ t0 att*λ, $Kan^r$ | This work |
| pMS16 | *oriR6K*, MCS $P_{fadH200}$-*lacZ t0 att*λ, $Kan^r$ | This work |
| pMK11 | *oriR6K*, MCS $P_{fadL200}$-*lacZ t0 att*λ, $Kan^r$ | This work |
| pMS15 | *oriR6K*, MCS $P_{fadM200}$-*lacZ t0 att*λ, $Kan^r$ | This work |
| pMK08 | *oriR6K*, MCS $P_{fadR200}$-*lacZ t0 att*λ, $Kan^r$ | This work |
| pLZ17 | *oriR6K*, MCS $P_{cyaR200}$-*lacZ t0 att*λ, $Kan^r$ | This work |
| pMK04 | *oriR6K*, MCS $P_{omrA200}$-*lacZ t0 att*λ, $Kan^r$ | This work |
| pLZ19 | *oriR6K*, MCS $P_{omrB200}$-*lacZ t0 att*λ, $Kan^r$ | This work |
| pLZ18 | *oriR6K*, MCS $P_{rprA200}$-*lacZ t0 att*λ, $Kan^r$ | This work |
| pMK10 | *oriR6K*, MCS $P_{cvpA350}$-*lacZ t0 att*λ, $Kan^r$ | This work |
| pMS07 | *oriR6K*, MCS $P_{ubiA350}$-*lacZ t0 att*λ, $Kan^r$ | This work |
| pLZ01 | *oriR6K*, MCS $P_{ubiB350}$-*lacZ t0 att*λ, $Kan^r$ | This work |
| pLZ08 | *oriR6K*, MCS $P_{ubiC1000}$-*lacZ t0 att*λ, $Kan^r$ | This work |
| pLZ12 | *oriR6K*, MCS $P_{ubiD600}$-*lacZ t0 att*λ, $Kan^r$ | This work |
| pLZ02 | *oriR6K*, MCS $P_{ubiE350}$-*lacZ t0 att*λ, $Kan^r$ | This work |
| pLZ05 | *oriR6K*, MCS $P_{ubiF400}$-*lacZ t0 att*λ, $Kan^r$ | This work |
| pLZ11 | *oriR6K*, MCS $P_{ubiG350}$-*lacZ t0 att*λ, $Kan^r$ | This work |
| pMS10 | *oriR6K*, MCS $P_{ubiH1000}$-*lacZ t0 att*λ, $Kan^r$ | This work |
| pLZ03 | *oriR6K*, MCS $P_{ubiI350}$-*lacZ t0 att*λ, $Kan^r$ | This work |
| pLZ06 | *oriR6K*, MCS $P_{ubiJ350}$-*lacZ t0 att*λ, $Kan^r$ | This work |
| pLZ07 | *oriR6K*, MCS $P_{ubiK350}$-*lacZ t0 att*λ, $Kan^r$ | This work |

| | | |
|---|---|---|
| pLZ10 | *oriR6K*, MCS $P_{ubiX350}$-*lacZ t0 att*λ, $Kan^r$ | This work |
| pMK08 | *oriR6K*, MCS $P_{ygfB500}$-*lacZ t0 att*λ, $Kan^r$ | This work |
| pZE12-*luc* | ColE1 *ori*, $P_{LlacO-1}$-*luc*, $Amp^r$ | Papenfort lab (16) |
| pMK05 (p*cpxQ*) | ColE1 *ori*, $P_{LlacO-1}$-*luc-cpxQ*, $Amp^r$ | This work |
| pNM46 | pBR322 carrying *lacI* and inducible $P_{lacO-1}$ promoter, $Amp^r$ | Storz lab (17) |
| pNV13 (p*omrA*) | pBR322 carrying *lacI* and $P_{lacO-1}$-*omrA*, $Amp^r$ | This work |
| pAM238 | pSC101 *ori*, inducible $P_{lac}$, $Spec^r$ | Collet lab (18) |
| pAM238-*nlpE* (p*nlpE*) | pSC101 *ori*, $P_{lac}$-*nlpE*, $Spec^r$ | Collet lab (18) |

**Table S4.** Primers used in this study

| Primer name | Sequence (5'→3') | Purpose | Source (Reference) |
|---|---|---|---|
| **Primers used to create pSH06, modified pSim5** | | | |
| SH67 | GGTGGTGAATTCTAAGAAATGTGCGCGGAACCC | Forward primer for amplifying β-lactamase gene with its promoter from pKD46 | This work |
| SH68 | GGTGGTCCATGGGGTCTGACAGTTACCAATGC | Reverse primer for amplifying β-lactamase gene with its promoter from pKD46 | This work |
| **Primers used to create sRNA deletion and SPA-tagged strains, and their verification** | | | |
| MS90 | TGAAGCTATTGAGTAGTAGCAACTCACGTTCCCAGTAGTAAACCCTGTTTATTCCGGGGATCCGTCGACC | Forward primer for making *cpxQ* deletion using Kan cassette from pKD13 | This work |
| MS91 | ATGCATTAAGCAGCAGGCAAATTGAGGATAAAAAAAACCCCCACAGCATGTGTAGGCTGGAGCTGCTTCG | Reverse primer for making *cpxQ* deletion using Kan cassette from pKD13 | This work |
| MK31 | TGAAGCTATTGAGTAGTAGCAACTCACGTTCCCAGTAGTAAACCCTGTTTCATGGGAATTAGCCATGGTCC | Forward primer for making *cpxQ* deletion using Cam cassette from pKD3 | This work |
| MK32 | ATGCATTAAGCAGCAGGCAAATTGAGGATAAAAAAAACCCCCACAGCATGTTGTGTAGGCTGGAGCTGCTTC | Reverse primer for making *cpxQ* deletion using Cam cassette from pKD3 | This work |
| MS92 | AGTCCGCAACCAAATGTATCG | Forward primer for the confirmation of *cpxQ* deletion | This work |
| MS93 | ATTTTAATCAGCAATAGCAGCG | Reverse primer for the confirmation of *cpxQ* deletion | This work |
| MS82 | GAAACCGATCACATACAGCTGCATTTATTAAGGTTATCATCCGTTTCGCTATTCCGGGGATCCGTCGACC | Forward primer for making *cyaR* deletion using Kan cassette from pKD13 | This work |
| MS83 | AAATCCTTTTATTTCATTGTATTACGCGTAAAAAATAAGCCCGTGTAAGGTGTAGGCTGGAGCTGCTTCG | Reverse primer for making *cyaR* deletion using Kan cassette from pKD13 | This work |
| MS94 | ATGGTTATACTGTGTGGCTCCC | Forward primer for the confirmation of *cyaR* deletion | This work |
| MS95 | TACAGCGATACATTGTGAAGC | Reverse primer for the confirmation of *cyaR* deletion | This work |
| MS86 | GCGTTTTCTCGCTGGCGAAGAGTCGTCGTGCAGACCACAATCAAGATCCCATTCCGGGGATCCGTCGACC | Forward primer for making *omrA* deletion using Kan cassette from pKD13 | This work |
| MS87 | ACGCGAGCGACAGTAAATTAGGTGCGAAAAAAAACCTGCGCATCCGCGCATGTAGGCTGGAGCTGCTTCG | Reverse primer for making *omrA* deletion using Kan cassette from pKD13 | This work |
| MS96 | TTGATAGGTGAAGTCAACTTCG | Forward primer for the confirmation of *omrA* deletion | This work |
| MS97 | AAAACACGGCACAAATTTCG | Reverse primer for the confirmation of *omrA* deletion | This work |
| MS88 | CGATTGACCGCTGGTGGCGTTTGGCTTCAGGTTGCTAAAGTGGTGATCCCATTCCGGGGATCCGTCGACC | Forward primer for making *omrB* deletion using Kan cassette from pKD13 | This work |

| | | | |
|---|---|---|---|
| MS89 | TCGGTTACTGTTACAGATTGATGACCGGCAAAAAAAACCTGCGCATCTGCTGTAGGCTGGAGCTGCTTCG | Reverse primer for making *omrB* deletion using Kan cassette from pKD13 | This work |
| MS98 | AAACGCTTACATTCTTTCAGTG | Forward primer for the confirmation of *omrB* deletion | This work |
| MS99 | AGAGCGTACCGAATAATCTCACC | Reverse primer for the confirmation of *omrB* deletion | This work |
| MS84 | AGACGAATCTGATCGACGCAAAAAGTCCGTATGCCTACTATTAGCTCACGATTCCGGGGATCCGTCGACC | Forward primer for making *rprA* deletion using Kan cassette from pKD13 | This work |
| MS85 | GCGAGGTAGCGAAGCGGAAAAATGTTAAAAAAAAGCCCATCGTGGGAGATGTAGGCTGGAGCTGCTTCG | Reverse primer for making *rprA* deletion using Kan cassette from pKD13 | This work |
| MS100 | ATAAGTAATTTCTCATCAGGCG | Forward primer for the confirmation of *rprA* deletion | This work |
| MS101 | AAGCGTTCATCGTGTAATGG | Reverse primer for the confirmation of *rprA* deletion | This work |
| MK27 | GGCACCGGATGCAGAAAAATCTGCCGGGTGATTTAGCCATTCAGGGGCGATCCATGGAAAAGAGAAGATGG | Forward primer to chromosomally SPA-tag FadR | This work |
| MK28 | AACAACAAAAAACCCCTCGTTTGAGGGGTTTGCTCTTTAAACGGAAGGGACATATGAATATCCTCCTTAG | Reverse primer to chromosomally SPA-tag FadR | This work |
| MK29 | GCGACTGGATCACGAAAGTG | Forward primer for the confirmation of FadR-SPA | This work |
| MK30 | CGCAAAGAAGTCCTGAAGCATG | Reverse primer for the confirmation of FadR-SPA | This work |
| SL38 | GCGACGAGAATTACGTGACGAAGCGCGCGGCAAAGTGGACAATAAAGCCTCCATGGAAAAGAGAAGATGG | Forward primer to chromosomally SPA-tag FadD | This work |
| SL39 | GCGTCAAAAAAAACGCCGGATTAACCGGCGTCTGACGACTGACTTAACGCCATATGAATATCCTCCTTAG | Reverse primer to chromosomally SPA-tag FadD | This work |
| SL40 | GAAGGATTCCTGCGCATTGTC | Forward primer for the confirmation of FadD-SPA | This work |
| SL41 | CTTCACACAAAGAAGCCAGCGC | Reverse primer for the confirmation of FadD-SPA | This work |
| SL34 | GAGTCTGAAGGTAAAGCCTGGCTGTTCGGTACTAACTTTAACTACGCGTTCTCCATGGAAAAGAGAAGATGG | Forward primer to chromosomally SPA-tag FadL | This work |
| SL35 | GAGTTAAAGTCACCTGCTATGCAGGTGACTTTATCCAGGCGAACGCGTTACATATGAATATCCTCCTTAG | Reverse primer to chromosomally SPA-tag FadL | This work |
| SL36 | GCCAATTCCTACCGCGACAG | Forward primer for the confirmation of FadL-SPA | This work |
| SL37 | CAGGTTTGTCTGTGGATACCG | Reverse primer for the confirmation of FadL-SPA | This work |
| SAK1 | GAGGCTATTCGGCTATGACTG | Forward primer specific to kanamycin cassette | (19) |
| SAK2 | TTCCATCCGAGTACGTGCTC | Reverse primer specific to kanamycin cassette | (19) |
| SH40 | CACCGTTGATATATCCCAATGGC | Forward primer specific to chloramphenicol cassette | This work |

| | | | |
|---|---|---|---|
| SH39A | TGATGAACCTGAATCGCCAG | Reverse primer specific to chloramphenicol cassette | This work |
| **Primers used for cloning and verification/sequencing of clones** | | | |
| MS107 | ACCGGGTACCAACGCGATCAAG TTCACTGC | Forward primer for cloning $P_{cpxP(-410/+223)}$ in pAH125 | This work |
| MS109 | CGTGAATTCCATTAACAGGAGGC TGTTCG | Reverse primer for cloning $P_{cpxP(-410/+223)}$ in pAH125 | This work |
| MS43 | ACCGGGTACCATTGATTTAAGAA TTTTCAGGTCGGATGC | Forward primer for cloning $P_{fadE1000}$ in pAH125 | (13) |
| MS129 | ACCGGGTACCGGCTTTAAAGCT GTCTGC | Forward primer for cloning $P_{fadE400}$ in pAH125 | This work |
| MS103 | ACCGGGTACCCATAAATGTAATA GACAAAATGC | Forward primer for cloning $P_{fadE200}$ in pAH125 | This work |
| MS44 | CGTGAATTCCCGTAGCGAGAATA CTCAAAATCATCAT | Reverse primer for cloning $P_{fadE1000}$/$P_{fadE400}$/$P_{fadE200}$ in pAH125 | (13) |
| MS102 | ACCGGGTACCATGATTTTTATAG AGCGAGG | Forward primer for cloning $P_{fadB200}$ in pAH125 | This work |
| MS51 | CGTGAATTCGGTGTCGCCTTTGT AAAGCAT | Reverse primer for cloning $P_{fadB200}$ in pAH125 | This work |
| MK44 | ACCGGGTACCATGGTTTTATGGC GGTCGTG | Forward primer for cloning $P_{fadD200}$ in pAH125 | This work |
| MK45 | CGTGAATTCGGTTAAGCCAAACC TTCTTCAA | Reverse primer for cloning $P_{fadD200}$ in pAH125 | This work |
| MS104 | ACCGGGTACCATAAGACGCGAC AAGCGTC | Forward primer for cloning $P_{fadH200}$ in pAH125 | This work |
| MS48 | CGTGAATTCCGAACAGCGACGG GTAGCTCAT | Reverse primer for cloning $P_{fadH200}$ in pAH125 | This work |
| MK42 | ACCGGGTACCAATTCACCCGAAT CCATGAGTG | Forward primer for cloning $P_{fadL200}$ in pAH125 | This work |
| MK43 | CGTGAATTCGTAAACAGGGTTTT CTGGCTCAT | Reverse primer for cloning $P_{fadL200}$ in pAH125 | This work |
| MS105 | ACCGGGTACCGTTGGCCTGAAG AAAGC | Forward primer for cloning $P_{fadM200}$ in pAH125 | This work |
| MS46 | CGTGAATTCCACGAACTTTGATT TGTGTTTGCAT | Reverse primer for cloning $P_{fadM200}$ in pAH125 | This work |
| MS106 | ACCGGGTACCCATGATGGTTTCC CTTAC | Forward primer for cloning $P_{fadR200}$ in pAH125 | This work |
| MS61 | CGTGAATTCCTTTGCGCCTTAAT GACCAT | Reverse primer for cloning $P_{fadR200}$ in pAH125 | This work |
| LZ51 | ACCGGGTACCTGGTTATACTGTG TGGCTCC | Forward primer for cloning $P_{cyaR200}$ in pAH125 | This work |
| LZ52 | CGTGAATTCGAGGTGGTTCCTG GTACAGC | Reverse primer for cloning $P_{cyaR200}$ in pAH125 | This work |
| MK01 | ACCGGGTACCGTTGAGCACATG AATTACACC | Forward primer for cloning $P_{omrA200}$ in pAH125 | This work |
| MK02 | CGTGAATTCCAATCAATACCTCT GGG | Reverse primer for cloning $P_{omrA200}$ in pAH125 | This work |
| LZ56 | ACCGGGTACCGGGTAAACGCTT ACATTC | Forward primer for cloning $P_{omrB200}$ in pAH125 | This work |
| MS123 | CGTGAATTCCACCTATCAATACC TCTGGG | Reverse primer for cloning $P_{omrB200}$ in pAH125 | This work |
| LZ53 | ACCGGGTACCAAATTCTCGAAGA ACTTGGC | Forward primer for cloning $P_{rprA200}$ in pAH125 | This work |
| LZ54 | CGTGAATTCTTCACTCAGGGGAT TTCC | Reverse primer for cloning $P_{rprA200}$ in pAH125 | This work |
| MS163 | ACCGGGTACCTGCCTCGAAAGA TAAGCTGAAAGG | Forward primer for cloning $P_{cvpA350}$ in pAH125 | This work |
| MS164 | CGTGAATTCTTATGGCGTAATCA ATCCAGACCAT | Reverse primer for cloning $P_{cvpA350}$ in pAH125 | This work |

| | | | |
|---|---|---|---|
| MS70 | ACCGGGTACCAGAATGAAATCCC CGAAGAACTG | Forward primer for cloning $P_{ubiA350}$ in pAH125 | This work |
| MS71 | CGTGAATTCATTCTGCGTCAGAC TCCACTCCAT | Reverse primer for cloning $P_{ubiA350}$ in pAH125 | This work |
| LZ1 | ACCGGGTACCAGCTTACCGCAC TGATTC | Forward primer for cloning $P_{ubiB350}$ in pAH125 | This work |
| LZ2 | CGTGAATTCACTTCACCTGGCGT CAT | Reverse primer for cloning $P_{ubiB350}$ in pAH125 | This work |
| LZ15 | ACCGGTCGACATGAAGATCTCGA TGGTTATATC | Forward primer for cloning $P_{ubiC1000}$ in pAH125 | This work |
| LZ16 | CGTGGTACCAACGCGGGGTGTG ACAT | Reverse primer for cloning $P_{ubiC1000}$ in pAH125 | This work |
| LZ17 | ACCGGGTACCAGATTAAGCAATA GCATGG | Forward primer for cloning $P_{ubiD600}$ in pAH125 | This work |
| LZ18 | CGTGAATTCATATTTCATGGCGT CCAT | Reverse primer for cloning $P_{ubiD600}$ in pAH125 | This work |
| LZ3 | ACCGGGTACCTCAAAGTCTCGAC AAAGC | Forward primer for cloning $P_{ubiE350}$ in pAH125 | This work |
| LZ4 | CGTGAATTCTTCTTGTGACTTATC CACCAT | Reverse primer for cloning $P_{ubiE350}$ in pAH125 | This work |
| LZ5 | ACCGGGTACCACGCCGATAATC AGGTCT | Forward primer for cloning $P_{ubiF400}$ in pAH125 | This work |
| LZ6 | CGTGAATTCCGTTGGTTGATTTG TCAT | Reverse primer for cloning $P_{ubiF400}$ in pAH125 | This work |
| LZ19 | ACCGGGTACCTTATAGGCTTTGT TCCAG | Forward primer for cloning $P_{ubiG350}$ in pAH125 | This work |
| LZ20 | CGTGAATTCGATTTTTCGGCATT CAT | Reverse primer for cloning $P_{ubiG350}$ in pAH125 | This work |
| MS72 | ACCGGGTACCATCAACTACTTAA CGGCCTGG | Forward primer for cloning $P_{ubiH1000}$ in pAH125 | This work |
| MS73 | CGTGAATTCCCGACGATGATTAC GCTCAT | Reverse primer for cloning $P_{ubiH1000}$ in pAH125 | This work |
| LZ7 | ACCGGGTACCAAACTCTGCACC CGATTG | Forward primer for cloning $P_{ubiI350}$ in pAH125 | This work |
| LZ8 | CGTGAATTCGCTACATCAACACT TTGCAT | Reverse primer for cloning $P_{ubiI350}$ in pAH125 | This work |
| LZ9 | ACCGGGTACCTTTGGTCTGCGTA ACGTC | Forward primer for cloning $P_{ubiJ350}$ in pAH125 | This work |
| LZ10 | CGTGAATTCCACTAAAGGTTTAA AAGGCAT | Reverse primer for cloning $P_{ubiJ350}$ in pAH125 | This work |
| LZ11 | ACCGGGTACCTTACCAGAATCAG GGCAG | Forward primer for cloning $P_{ubiK350}$ in pAH125 | This work |
| LZ12 | CGTGAATTCTTTTTCGGGTCAAT CAT | Reverse primer for cloning $P_{ubiK350}$ in pAH125 | This work |
| LZ13 | ACCGGGTACCAAGTTGATGAAAT TCGCC | Forward primer for cloning $P_{ubiX350}$ in pAH125 | This work |
| LZ14 | CGTGAATTCCCTACAATGAGTCG TTTCAT | Reverse primer for cloning $P_{ubiX350}$ in pAH125 | This work |
| NV07 | ACCGGAATTCTTGGTTAGTTTTTT CGGTGATGCG | Forward primer for cloning $P_{ygfB500}$ in pAH125 | This work |
| NV08 | CGTGAATTCAGGCATTTCGTTCT GTATAGACAT | Reverse primer for cloning $P_{ygfB500}$ in pAH125 | This work |
| BS106 | TTGTCGGTGAACGCTCTCCT | Forward primer for sequencing/verification of clones in pAH125 | (13) |
| MS49 | TAAAACGACGGCCAGTGAATCC | Reverse primer for sequencing/verification of clones in pAH125 | (13) |
| MS133 | ACCGTCTAGATTTTCCTTGCCAT AGACACC | Forward primer for cloning *cpxQ* in pZE12-*luc* | This work |
| MS134 | CGTTCTAGAAACTGACGCTAGTA TAACGG | Reverse primer for cloning *cpxQ* in pZE12-*luc* | This work |

| | | | |
|---|---|---|---|
| MS135 | TGTTTGTGGACGAAGTACC | Forward primer for sequencing/verification of clones in pZE12-*luc* | This work |
| MS136 | TTTGTCCTACTCAGGAGAGC | Reverse primer for sequencing/verification of clones in pZE12-*luc* | This work |
| MK53 | ACCTGACGTCCCCAGAGGTATTGATTGGTGAGATT | Forward primer for cloning *omrA* in pNM46 | This work |
| MK54 | CGTGAATTCGAGCGACAGTAAATTAGGTGCG | Reverse primer for cloning *omrA* in pNM46 | This work |
| MK55 | GAATAAGGGCGACACGGAAATG | Forward primer for sequencing/verification of clones in pNM46 | This work |
| MK56 | CACTTTATGCTTCCGGCATTG | Reverse primer for sequencing/verification of clones in pNM46 | This work |
| **Primers used for strain verification of single integrants at the *att*λ site** | | | |
| GA22 | GGCATCACGGCAATATAC | Forward primer specific to *att*λ site on *E. coli* chromosome for confirmation of single-copy integration of reporter plasmids | (20) |
| GA23 | ACTTAACGGCTGACATGG | Forward primer specific to pAH125 to identify multiple-copy integration of reporter plasmids | (20) |
| GA25 | TCTGGTCTGGTAGCAATG | Reverse primer specific to *att*λ site on *E. coli* chromosome for confirmation of single-copy integration of reporter plasmids | (20) |
| GA29 | TGCGAGGCTTTGTGCTTC | Reverse primer specific to pAH125 to identify multiple-copy integration of reporter plasmids | (20) |
| **Primers used for qRT-PCR** | | | |
| MK07A | CCTTGCCATAGACACCATCC | Forward primer for *cpxQ* | This work |
| MK08B | ACAGCATGTGGGGGAAGAC | Reverse primer for *cpxQ* | This work |
| MS157 | ACCCATAAAATGCTAGCTGTACCAGG | Forward primer for *cyaR* | This work |
| MS158 | GTAAGGGAGATTACACAGGCTAAGGAGG | Reverse primer for *cyaR* | This work |
| MS151 | GGTATTGATTGGTGAGATTATTCGG | Forward primer for *omrA* | This work |
| MS152 | TTGGTGCAAGAGACAGGGTACG | Reverse primer for *omrA* | This work |
| MS159 | CCAGAGGTATTGATAGGTGAAGTC | Forward primer for *omrB* | This work |
| MS160 | CAGGCTGGTGTAATTCATGTGC | Reverse primer for *omrB* | This work |
| MS161 | AAGCATGGAAATCCCCTGAGTG | Forward primer for *rprA* | This work |
| MS162 | ATCGTGGGAGATGGCAAAGACTAC | Reverse primer for *rprA* | This work |
| MS33 | GAATTTTTGTCCCTGTTCCTCG | Forward primer for *fadB* | This work |
| MS34 | ATGATGCCCAGTTTGGTTTCC | Reverse primer for *fadB* | This work |
| MS145 | CCTAATTTATTGCAATATCCG | Forward primer for *fadD* | This work |
| MS146 | GTGTGAGCAAAGTTAGACACG | Reverse primer for *fadD* | This work |
| KJ87 | CTGACCTGGAACAAACGCTACATTAC | Forward primer for *fadE* | (21) |
| KJ88 | TACGTTCAGCGGGAAGTGG | Reverse primer for *fadE* | (21) |
| MS35 | AAATTTTGCATACCGGGC | Forward primer for *fadH* | This work |
| MS36 | ATCACCTCTACACCGTCGTATC | Reverse primer for *fadH* | This work |

| | | | |
|---|---|---|---|
| KJ36 | ACTTTGTTGCACCGATTAACG | Forward primer for *fadL* | (21) |
| KJ37 | ATACGCACCGCTTAAGTTCAGG | Reverse primer for *fadL* | (21) |
| MS39 | ACAGTTTTCAGTGGATGACGGC | Forward primer for *fadM* | This work |
| MS40 | TGTAATGACCTGGCTTAAGATGC | Reverse primer for *fadM* | This work |
| KJ18 | TGACGCTGAATACGATGACC | Forward primer for *ubiA* | This work |
| KJ19 | TCAATGGCACCGACTCACTC | Reverse primer for *ubiA* | This work |
| KJ20 | TTGAAATCAAGCCGCTGG | Forward primer for *ubiB* | This work |
| KJ21 | AGCGAGCCAGACGGTAGATAAG | Reverse primer for *ubiB* | This work |
| KJ22 | GAACAGCAGGGAAAAACGG | Forward primer for *ubiC* | This work |
| KJ23 | ACAGGAACGACGGTACGACC | Reverse primer for *ubiC* | This work |
| KJ24 | AATCGCATTCCCATTATGACC | Forward primer for *ubiD* | This work |
| KJ25 | GACAGCCAGCGCATAATCAG | Reverse primer for *ubiD* | This work |
| KJ26 | TATCGGTGTGATTGGCAACG | Forward primer for *ubiE* | This work |
| KJ27 | AGCACGCGATACATTGAACG | Reverse primer for *ubiE* | This work |
| KJ28 | CAGGAGCTGGAGCTGAAAGG | Forward primer for *ubiF* | This work |
| KJ29 | ATCGTTCTCGCACTGGACG | Reverse primer for *ubiF* | This work |
| KJ30 | TGCTCGATGTCGGTTGTGG | Forward primer for *ubiG* | This work |
| KJ31 | GTTTCCTGCACGTAATCCACC | Reverse primer for *ubiG* | This work |
| KJ32 | TTGATGGACGAGCGATAGC | Forward primer for *ubiH* | This work |
| KJ33 | TAATCTTCTGCGGCGAGG | Reverse primer for *ubiH* | This work |
| KJ34 | CGATCAAAGCATGGGCTATAG | Forward primer for *ubiI* | This work |
| KJ35 | ATCTTTCAGCGTCAGGAAGG | Reverse primer for *ubiI* | This work |
| MK05 | AGGAATTGAAAGTCTGCTC | Forward primer for *ubiJ* | This work |
| MK06 | CATTCGCCCAGTACATCAAC | Reverse primer for *ubiJ* | This work |
| KJ38 | AGGTTCACGAATCAATGCCTAAA G | Forward primer for *ubiK* | This work |
| KJ39 | AGCAGCGCCAGTTTTTCAC | Reverse primer for *ubiK* | This work |
| MK03 | AGACCTTATCCCTCGAAACG | Forward primer for *ubiX* | This work |
| MK04 | TATAGCTATGGACAATGCCG | Reverse primer for *ubiX* | This work |
| MS171 | TACACCGGTGAAAAATCCACACA GG | Forward primer for *icd* | This work |
| MS172 | TTGGAGTGCCCTGATAGTAACGT ACC | Reverse primer for *icd* | This work |
| MS175 | GAAATTACGCGGTTCAGTCAATC C | Forward primer for *aceA* | This work |
| MS176 | CATCCCGACAGATAGACTGCTTC A | Reverse primer for *aceA* | This work |
| MS173 | GGATCCAGTATCAGGGATTCAAA ACC | Forward primer for *maeA* | This work |
| MS174 | AACGGCGGTAGATCTCAGAAAAA CG | Reverse primer for *maeA* | This work |
| MS197 | GATGTTCTCCATTGATACCGAAT CC | Forward primer for *ppsA* | This work |
| MS198 | ACCATGCGGATTTTTTTCGACC | Reverse primer for *ppsA* | This work |
| MS179 | TCGTTAATCAGGTAAAAGACGCT GG | Forward primer for *nuoF* | This work |
| MS180 | TCCATCAACAGGCGGTCTTTATA GG | Reverse primer for *nuoF* | This work |
| MS177 | AAACGCGAATGTACCCGGAAGA G | Forward primer for *nuoI* | This work |

| | | | |
|---|---|---|---|
| MS178 | GTGAGAAGTTGATGCGGAAAAATTCC | Reverse primer for *nuoI* | This work |
| MS187 | AAAGGACAGATTGGTCTGGAGCAACG | Forward primer for *cyoA* | This work |
| MS188 | TCCAGACCACAGCTTCCACTTTATTG | Reverse primer for *cyoA* | This work |
| MS189 | TCACCACTACGATCAGATCTTTACCG | Forward primer for *cyoB* | This work |
| MS190 | GTTAACCAGAATCACACCAACAACG | Reverse primer for *cyoB* | This work |
| MS193 | AAAGTGATTGGTCACCTTGCACACG | Forward primer for *atpG* | This work |
| MS194 | TTGTCGGTCCAGGTCTTCATTTCC | Reverse primer for *atpG* | This work |
| MS195 | TAACCAAAAACGAACAAATGGCAGAG | Forward primer for *atpH* | This work |
| MS196 | GAATAAACTGCTCCAGAACATCCGG | Reverse primer for *atpH* | This work |

*Restriction sites are underlined

**Dataset S1 (separate .xlsx file).** DESeq output of genes in various comparison groups namely, WT_TBK-Ole vs. WT_TBK-Brij, Δ*cpxR*_TBK-Brij vs. WT_TBK-Brij, and Δ*cpxR*_TBK-Ole vs. WT_TBK-Ole, listed in three separate sheets. Base mean refers to the average of the normalized count values, divided by size factors, taken over all samples. log2FoldChange is the effect size estimate indicating how much the gene's expression seems to have changed between the comparison and control groups. Fold change refers to the ratio of gene expression measured between the two comparison groups. The lfcSE is the standard error estimate for the log2 fold change estimate. The stat is the value of the test statistic for the gene or transcript. The pvalue column represents the p-value of the test for the gene or transcript and padj is the adjusted p-value for multiple testing for the gene or transcript.

**Dataset S2 (separate .xlsx file).** Gene Set Enrichment Analysis of pathways in WT and Δ*cpxR* cultured in TBK-Brij and TBK-Ole. Statistical significance of the enrichment of pathways for various comparison groups i.e., WT_TBK-Ole vs. WT_TBK-Brij, Δ*cpxR*_TBK-Brij vs. WT_TBK-Brij and Δ*cpxR*_TBK-Ole vs. WT_TBK-Ole. Name is the pathway name as reported by EcoCyc. Size is the number of genes within this group. The Normalized Enrichment Score (NES) reflects the maximal enrichment of a pathway, normalized for set size and average enrichment across the dataset. NES is the primary metric for comparing significance between gene sets. The Nominal (NOM) p-value reflects the statistical significance of a given NES score without multiple hypothesis correction. The False Discovery Rate (FDR) q-value reflects the fraction of gene sets with a given NES score that are expected to be false positives. Pathways related to LCFA metabolism and ubiquinone biosynthesis are highlighted in light blue. Three different comparison groups are listed in three separate sheets.
